# START domain mediates *Arabidopsis* GLABRA2 transcription factor dimerization and turnover independently of homeodomain DNA binding

**DOI:** 10.1101/2021.11.23.469727

**Authors:** Thiya Mukherjee, Bibek Subedi, Aashima Khosla, Erika M. Begler, Preston M. Stephens, Adara L. Warner, Ruben Lerma-Reyes, Kyle A. Thompson, Sumedha Gunewardena, Kathrin Schrick

**Affiliations:** Division of Biology, Kansas State University, Manhattan, KS 66506, USA; Molecular, Cellular and Developmental Biology, Kansas State University, Manhattan, KS 66506, USA; Donald Danforth Plant Science Center, Olivette, MO 63132, USA; Department of Botany and Plant Sciences, University of California, Riverside, CA 92521, USA; Interdepartmental Genetics, Kansas State University, Manhattan, KS 66506, USA; Department of Molecular and Integrative Physiology, University of Kansas Medical Center, Kansas City, KS 66160, USA

## Abstract

Class IV homeodomain leucine-zipper transcription factors (HD-Zip IV TFs) are key regulators of epidermal differentiation that are characterized by a DNA-binding homeodomain (HD) in conjunction with a lipid-binding domain termed START (<u>St</u>eroidogenic <u>A</u>cute <u>R</u>egulatory (StAR)-related lipid <u>T</u>ransfer). Previous work established that the START domain of GLABRA2 (GL2), a HD-Zip IV member from *Arabidopsis*, is required for transcription factor activity. Here, we address the functions and possible interactions of START and the HD in DNA binding, dimerization, and protein turnover. Deletion analysis of the HD and missense mutations of a conserved lysine (K146) result in phenotypic defects in leaf trichomes, root hairs and seed mucilage, similar to those observed for START domain mutants, despite nuclear localization of the respective proteins. *In vitro* and *in vivo* experiments demonstrate that while HD mutations impair binding to target DNA, the START domain is dispensable for DNA binding. Vice versa, protein interaction assays reveal impaired GL2 dimerization for multiple alleles of START mutants, but not HD mutants. Using *in vivo* cycloheximide chase experiments, we provide evidence for the role of START, but not HD, in maintaining protein stability. This work advances our mechanistic understanding of HD-Zip TFs as multidomain regulators of epidermal development in plants.

## INTRODUCTION

Class IV homeodomain leucine-zipper transcription factors (HD-Zip IV TFs) are critical regulators of epidermal and sub-epidermal differentiation in plants (Nakamura et al., 2006). In *Arabidopsis*, there are 16 START domain-containing transcription factors belonging to the HD-Zip IV family. The founding member, GLABRA2 (GL2), functions in a complex regulatory network to dictate epidermal cell fate. Genetic studies initially showed that GL2 promotes trichome differentiation in shoots (Rerie et al., 1994) and mucilage biosynthesis in seeds (Western et al., 2001; Western et al., 2004), while it suppresses hair formation in non-hair root cells (Di Cristina et al., 1996; Masucci et al., 1996). More recent mutant analyses revealed that GL2 regulates the accumulation of seed oil (Shen et al., 2006; Shi et al., 2012) and anthocyanin biosynthesis (Wang et al., 2015). In addition to GL2, the functionally redundant family members *Arabidopsis thaliana* MERISTEM LAYER1 (ATML1) and PROTODERMAL FACTOR2 (PDF2) have been extensively studied for their developmental role in specification of the epidermis in the embryo (Abe et al., 2003; Ogawa et al., 2015; Iida and Takada, 2021). HD-Zip IV proteins are comprised of four defined domains including a DNA-binding homeodomain (HD), a leucine-zipper domain termed Zipper Loop Zipper (ZLZ), a <u>St</u>eroidogenic <u>A</u>cute <u>R</u>egulatory (<u>StAR</u>) protein-related lipid <u>T</u>ransfer (START) domain, and a START-associated domain (SAD or STAD) (Schena and Davis, 1994; Di Cristina et al., 1996; Ponting and Aravind, 1999; Riechmann et al., 2000; Schrick et al., 2004).

The HD is a DNA-binding domain of ~60 amino acids that is highly conserved in the eukaryotes (Carrasco et al., 1984; McGinnis et al., 1984; Gehring et al., 1990; Bürglin, 1994; Derelle et al., 2007). It was discovered in *Drosophila* genes, whose mutation or overexpression cause homeotic transformations affecting the body pattern (Garber et al., 1983; McGinnis et al., 1984; Scott and Weiner, 1984). The 3D structure of the HD shows a flexible amino-terminus and three α-helices, the second and third of which form a helix-turn-helix motif (Gehring et al., 1990; Kissinger et al., 1990; Liu et al., 1990; Otting et al., 1990; Gehring, 1994). DNA binding specificity occurs through major groove contacts via the third helix as well as minor groove contacts via the amino-terminal region (Schofield, 1987; Mann et al., 2009; Burglin and Affolter, 2016). The first reported homeobox gene from plants was maize *Knotted-1* (*Kn-1*), whose gain-of-function phenotype is marked by uncontrolled cell divisions in lateral veins, forming outpocketings or knots (Vollbrecht et al., 1991). Since the discovery of *Kn-1*, numerous plant homeobox genes have been identified and implicated in a wide variety of developmental processes. In *Arabidopsis*, there are 110 predicted HD proteins that are subdivided into 14 classes including HD-Zip TFs (Classes I-IV) (Mukherjee et al., 2009).

HD-Zip III and IV TFs contain a START domain that occurs in the middle of the protein, carboxy-terminal to the HD-Zip. Initially discovered in mammalian StAR (<u>St</u>eroidogenic <u>A</u>cute <u>R</u>egulatory) cholesterol-binding transporters (Kallen et al., 1998), START (StAR-related lipid Transfer) domains are modules of ~210 amino acids (Ponting and Aravind, 1999) conserved across plants and animals as well as several protists, bacteria and archaea (Iyer et al., 2001). The domain structure comprises of an α/β helix-grip fold that forms a binding pocket (Tsujishita and Hurley, 2000; Thorsell et al., 2011). Human START domain-containing proteins bind ligands such as sterols, phospholipids, ceramides, bile acids and fatty acids (Kallen et al., 1998; Tsujishita and Hurley, 2000; Roderick et al., 2002; Kudo et al., 2008; Létourneau et al., 2012; Tillman et al., 2020). In mammals, START domains function in lipid/sterol transport or metabolism and several act in intracellular signaling pathways (Clark and Stocco, 1995; Lin et al., 1995; Watari et al., 1997; Stocco, 2001; Soccio and Breslow, 2003).

Studies in plants indicate similar roles for START domains in lipid transfer and/or lipid sensing. In the liverwort *Marchantia polymorpha*, a START domain-containing protein (STAR2) was recently implicated in the incorporation of ER-derived C20 fatty acids into chloroplast glycolipids (Hirashima et al., 2021). Our previous work showed that the START domain is essential for transcription factor activity of HD-Zip IV TF GL2 (Schrick et al., 2014). Several mechanisms have been proposed to explain how the ligand-bound START domain can stimulate transcription factor activity, including conformational changes, stabilization of protein levels, and/or promotion of protein–protein interactions (Schrick et al., 2014). However, lack of detailed information on how START modulates transcription factor activity poses as a major challenge.

The goal of this study was to understand how the START domain modulates HD-Zip IV transcription factor activity either independently or in synergy with the HD. We generated HD mutations in wild-type and START domain-deleted backgrounds and studied their effect on transcription factor function. HD mutants lacking the predicted DNA-binding α-helices or containing missense mutations in a conserved lysine imparted defective phenotypes in trichomes, roots and seeds, similar to those of START mutants. While HD mutations abolished DNA binding, multiple mutations affecting the START domain retained DNA binding affinity *in vitro* and *in vivo*. Gel shift experiments led to the surprising discovery that HD-Zip IV proteins lacking the START domain can bind DNA as monomers. Using interaction studies in yeast and *Nicotiana benthamiana* we confirmed dimerization defects from mutations affecting START but not HD. Finally, cycloheximide chase experiments showed that protein levels drop rapidly in several START domain mutants but not in HD mutants, consistent with a unique role of START in maintaining protein stability.

## RESULTS

### HD and START domain are highly conserved across the HD-Zip IV TFs

The HD is evolutionarily conserved across all major eukaryotic lineages (Carrasco et al., 1984; McGinnis et al., 1984; Derelle et al., 2007; Mukherjee et al., 2009). We found that the crystal structure of HOMEOBOX PROTEIN ENGRAILED-2 (Gg-En-2) from chicken exhibits the closest homology with the HD from HD-Zip IV members. Aligning HDs from all 16 HD-Zip IV TFs (GL2, PDF2, ANL2, ATML1, HDG1-12) with reference to Gg-En-2 revealed highly conserved α-helices as well as amino acid residues (**Supplemental Figure 1A**). The lysine at position 45 of the HD displays 100% conservation in HD-Zip IV members (**Figure 1A; Supplemental Figure 1A**). This conserved lysine selectively recognizes the 3’ CC dinucleotide adjacent to the TAAT core sequence (Amendt et al., 1998; Cazorla et al., 2000; Lebel et al., 2001) in the vertebrate Bicoid-like transcription factors PITX2 and PITX3 (Semina et al., 1996; Gage and Camper, 1997; Semina et al., 1997; Smidt et al., 1997).

**Figure 1.**
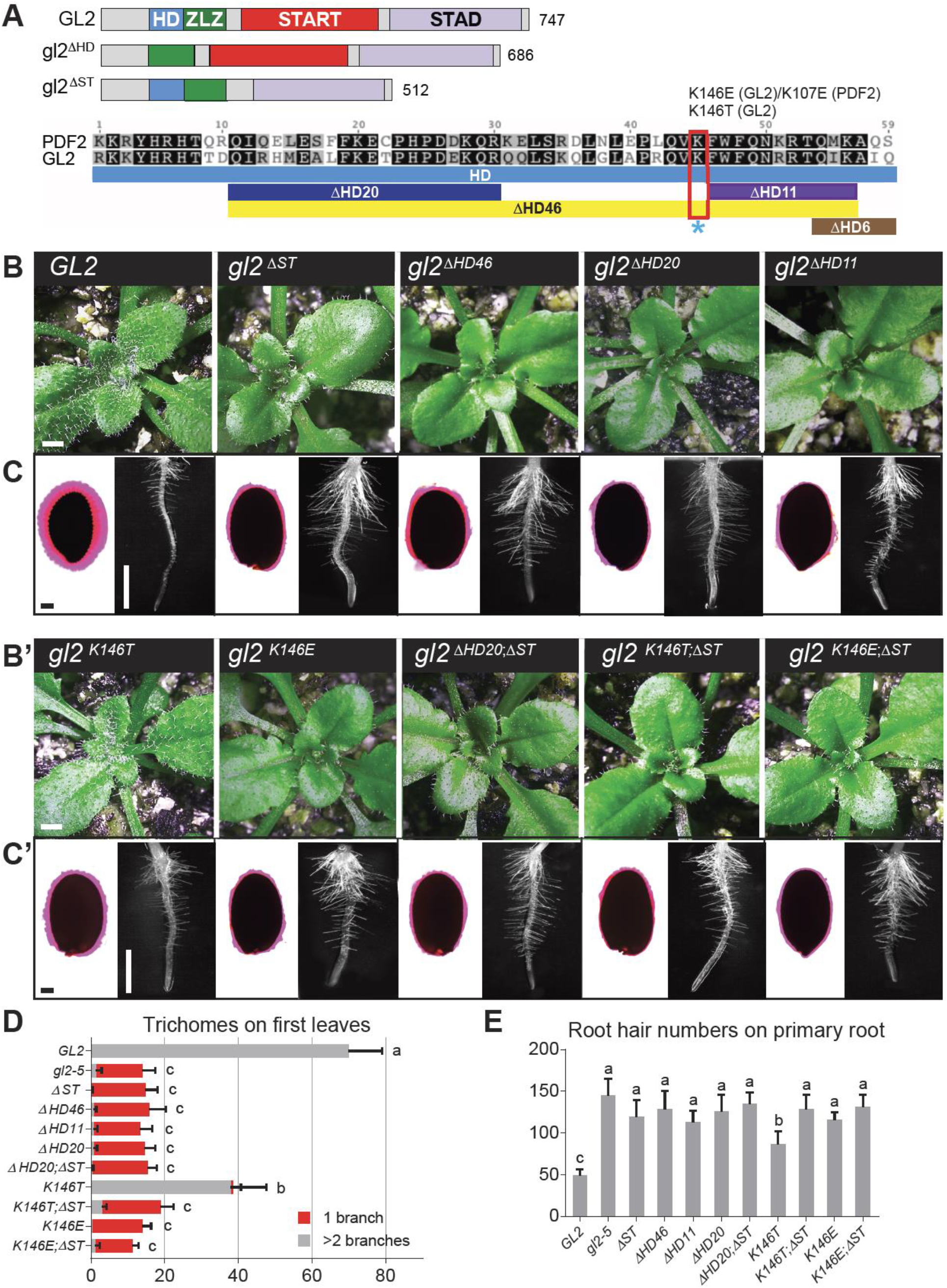
HD mutants exhibit loss-of-function phenotype in leaves, seeds, and roots. **(A)** Schematic of wild-type GL2 and mutant gl2^ΔHD^ and gl2^ΔSTART^ proteins containing homeodomain (HD), Zipper-Loop-Zipper (ZLZ), Steroidogenic Acute regulatory protein-Related lipid Transfer (START) and START-Associated Domain (STAD). Amino acid lengths of proteins are indicated (right). Alignment of GL2 and PDF2 HD domains generated by Geneious 7.0.6 (Biomatters). Identical and similar amino acid residues are indicated by black and gray shading, respectively. See **Supplemental Figure 1A**. HD deletions are shown as blue, purple, yellow, or brown tracts. Lysine targeted for missense mutations (K-E or K-T) is indicated by red box and blue asterisk. **(B)** and **(B’)** Wild-type *GL2* rosettes display trichomes covering leaf surfaces. In contrast, mutants (*gl2^ΔST^, gl2^ΔHD46^, gl2^ΔHD20^, gl2^ΔHD11^, gl2^K146E^, gl2^ΔHD20;ΔST^, gl2^K146T;ΔST^, gl2^K146E;ΔST^*) exhibit trichome differentiation defects. *gl2^K146T^* displays an intermediate phenotype. Bar = 1 mm. **(C)** and **(C’)** Left: Ruthenium red staining of seeds to assay mucilage production. Wild-type *GL2* seeds display a normal mucilage layer, while the various *gl2* mutants exhibit defective mucilage production. *gl2^K146T^* displays a partial phenotype. Bar = 100 μm. Right: Root hair phenotypes of 4-5 day-old seedlings. In contrast to wild-type *GL2* and the partial phenotype of *gl2^K146T^*, all other mutants exhibit excessive root hair formation. Bar = 1 mm. **(D)** Total number and branching pattern of trichomes on the first leaves. Gray bars indicate trichomes with 2-5 branches in plants expressing wild-type EYFP-tagged GL2. Mutant plants of the genotypes *gl2-5*, and the other *gl2* mutants exhibit unbranched trichomes, indicated by red bars. *gl2^K146T^* plants display a partial trichome differentiation defects. Values are mean ± SD for n = 20 plants. Letters denote significant differences from ordinary one-way ANOVA (p < 0.05) using Tukey’s multiple comparisons test. **(E)** Numbers of root hairs on primary roots of 4-5 day-old seedlings. Genotypes correspond to those in **(D)**. All the *gl2* mutants display increases in the number of root hairs. The *gl2^K146T^* seedlings exhibit an intermediate phenotype. Values are mean ± SD for n = 6-10 seedlings. Letters denote significant differences from one-way ANOVA (p < 0.05) using Tukey’s multiple comparisons test.

The conserved features of the START domain among HD-Zip IV members were studied using a structure-based sequence alignment with STARD5, a mammalian START protein with known crystal structure (Soccio et al., 2002; Rodriguez-Agudo et al., 2005; Thorsell et al., 2011). Despite low sequence identity, the alignment revealed consensus α helices and β strands, as well as conserved amino acids implicated in ligand binding (**Supplemental Figure 1B**).

To study the role of the HD in HD-Zip IV TFs, we generated several constructs with HD mutations in both wild-type GL2 (and/or PDF2) and START-deleted (ΔST) backgrounds (**Figure 1A**). The gl2^ΔST^ mutant lacks 235 residues encoding the START domain (Schrick et al., 2014). We constructed HD deletion mutants that remove the consensus α-helices in GL2 (ΔHD11, ΔHD20, ΔHD46) and missense mutants (K-E, K-T) targeting a highly conserved lysine (K146 in GL2; K107 in PDF2) (**Figure 1A**). This positively-charged lysine was replaced with a negatively-charged glutamic acid or with a neutral polar residue, threonine. Transgenic lines were constructed in the *gl2-5* null mutant background with wild-type *GL2* or mutant *gl2* expressed under the native *GL2* promoter with the enhanced yellow fluorescent protein (EYFP) translationally fused at the amino-terminus. For each genotype, we screened >20 independent T1 transformants to monitor nuclear expression of EYFP and phenotypic complementation of *gl2-5* (**Supplemental Table 1**), and chose representative homozygous T3 lines for further characterization.

### Mutations affecting the HD of GL2 exhibit defects in leaf, root and seed epidermis

Deletion of the START domain in *GL2* was previously shown to cause visible defects in differentiation of the epidermis in the leaf (trichomes), root (non-hair cells), and seed (seed coat mucilage) similar to *gl2-5* null mutant phenotypes (Schrick et al., 2014). We examined the phenotypic effects of mutations in the HD in presence or absence of START domain deletion (**Figure 1**). Unlike wild-type plants, rosette leaves from HD mutants lacking consensus α-helices (*gl2^ΔHD11^, gl2^ΔHD20^, gl2^ΔHD46^*) or containing a K-E substitution in the conserved lysine (*gl2^K146E^*) displayed trichome defects and were indistinguishable from ΔST or *gl2-5* null mutants (**Figure 1B and 1B’**). Another missense mutation in the same lysine (K146T) resulted in partial rescue (**Figure 1B’**). Importantly, three lines harboring both HD and START domain mutations (gl2^ΔHD20;ΔST^, gl2^K146E;ΔST^, gl2^K146T;ΔST^) were also defective in trichome differentiation similar to ΔST mutants (**Figure 2A’**). Quantification of trichomes on the first leaves revealed a reduction in total number as well as the abnormal presence of unbranched trichome in these mutants (**Figure 1D**).

**Figure 2.**
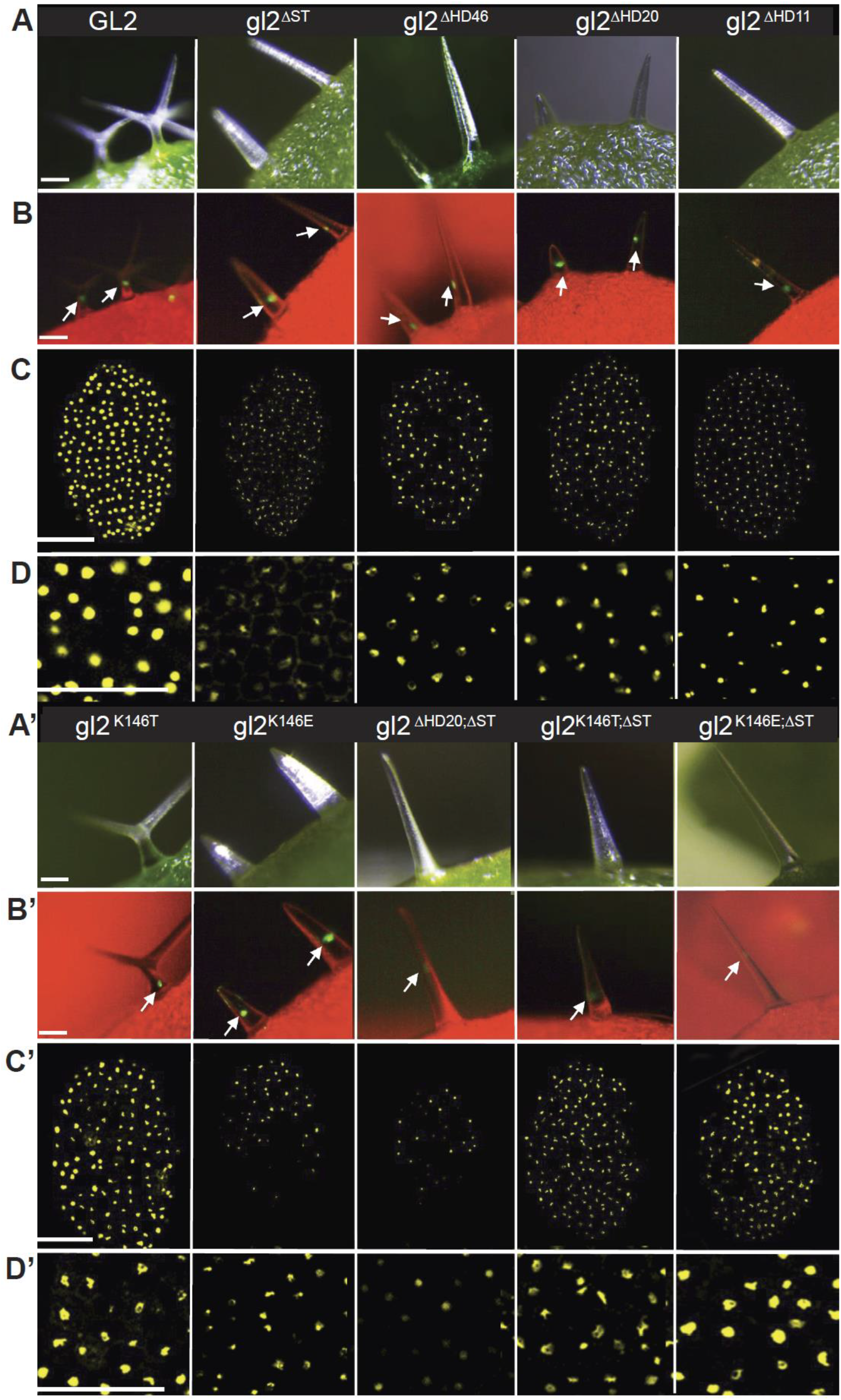
Nuclear localization of wild-type EYFP-tagged GL2 and mutant gl2 transcription factors. **(A)** and **(A’)** Imaging of leaf trichomes under white light (green, chlorophyll). Bar = 100 μm. **(B)** and **(B’)** Matching fluorescence images (red, chlorophyll) indicate nuclear localization of EYFP-tagged GL2, gl2^ΔST^, gl2^ΔHD46^, gl2^ΔHD20^, gl2^ΔHD11^, gl2^K146T^, gl2^K146E^, gl2^ΔHD20;ΔST^, gl2^K146T;ΔST^ and gl2^K146E;ΔST^ proteins in trichome cells. Arrows mark the brightest nuclear signal in each image. Bar = 100 μm. **(C)** and **(C’)** Confocal laser scanning microscopy of developing seeds reveals nuclear expression of EYFP-tagged wild-type and mutant proteins in seed coat cells. Bar = 100 μm. **(D)** and **(D’)** Magnified images from **(C)** and **(C’)**. Bar = 50 μm.

Apart from trichome defects, mutation of HD resulted in excessive root hair formation characteristic of *gl2-5* null mutants, irrespective of accompanying START domain mutation (**Figures 1C, 1C’ and 1E**). To assess the effect of mutations on differentiation of the mucilage secretory cells, imbibed seeds were stained with ruthenium red, a dye that binds to pectinaceous polysaccharides. Reduced seed coat mucilage was observed in HD mutants gl2^ΔHD11^, gl2^ΔHD20^, gl2^ΔHD46^, and gl2^K146E^, while the K146T mutation resulted in a partial mucilage defect (**Figures 1C and 1C’**) consistent with partial phenotypes observed in trichomes and roots (**Figures 1D and 1E**). HD and START domain double mutants (gl2^ΔHD20;ΔST^, gl2^K146E;ΔST^, gl2^K146T;ΔST^) were also defective in seed mucilage similar to ΔST mutants (**Figures 1C and 1C’**). These results demonstrate that both the HD and START domain are critical for epidermal differentiation in multiple cell types, and that mutation in one domain is not able to suppress the loss-of-function phenotype in the other domain.

### HD and START domain mutants retain nuclear localization of GL2

GL2 is expressed in developing trichomes of the leaf epidermis (Rerie et al., 1994; Szymanski et al., 1998), non-hair cell files in the root epidermis (Di Cristina et al., 1996; Masucci et al., 1996) and hypocotyl (Hung et al., 1998), and seed coat cells (Western et al., 2004). As a transcription factor, GL2 is presumably translocated to the nucleus where it binds DNA to regulate gene expression. Using EYFP fusions to GL2, it was previously shown that the START domain is not necessary for nuclear localization (Schrick et al., 2014). Here we monitored the expression of *EYFP:GL2* transgenes harboring various combinations of HD and START domain mutations in leaf trichomes and developing seeds. Using live imaging of trichomes, we observed nuclear localization of the EYFP-tagged proteins in all the HD mutants similar to that for wild-type GL2 and gl2^ΔST^ (**Figures 2A, 2A’, 2B and 2B’**). Nuclear localization remained intact in trichomes of mutants for both the HD and START domains (gl2^ΔHD20;ΔST^, gl2^K146E;ΔST^, gl2^K146T;ΔST^) (**Figures 2A’ and 2B’**). Confocal laser scanning microscopy of developing seeds indicated nuclear localization of HD and START mutant proteins in the seed coat cells (**Figures 2C, 2C’, 2D and 2D’**). The results demonstrate that both HD and START domain are dispensable for translocation of GL2 to the nucleus.

### START domain missense mutants exhibit loss-of-function despite nuclear localization of GL2

We next tested whether missense mutations in the START domain result in phenotypic defects similar to those seen in the START deletion mutant, *gl2^ΔST^* (**Figure 3A**). Transgenic lines carrying *EYFP:gl2* constructs were established according to the strategy described above for the HD mutants and *gl2^ΔST^*. Phenotypic analysis showed that three START missense mutants (*gl2^E375G;R392M^, gl2^Amo^, gl2^L480P^*) and a mutant deleted for the six terminal amino acids of the HD and ZLZ domain (*gl2*^Δ*HD6*;Δ*ZLZ*^), display leaf trichome defects and seed mucilage defects (**Figure 3B, 3C and 3C’**), as well as excess root hair formation (**Supplemental Figure 2A**) similar to the *gl2-5* null mutant. The mutations in *gl2^E375G;R392M^* disrupt a salt bridge that is thought to be critical for ligand binding (Roostaee et al., 2009). In *gl2^Amo^*, the R386L and E387L missense mutations, which remove charged residues, are followed by an 11-residue duplication in a predicted ligand-contact region (**Figure 3A and Supplemental Figure 1B**). In *gl2^L480P^*, the mutation affects a lipid contact site in the C-terminal α-helix of the START domain (Roostaee et al., 2009; Alpy and Tomasetto, 2014; Wojciechowska, 2021).

**Figure 3.**
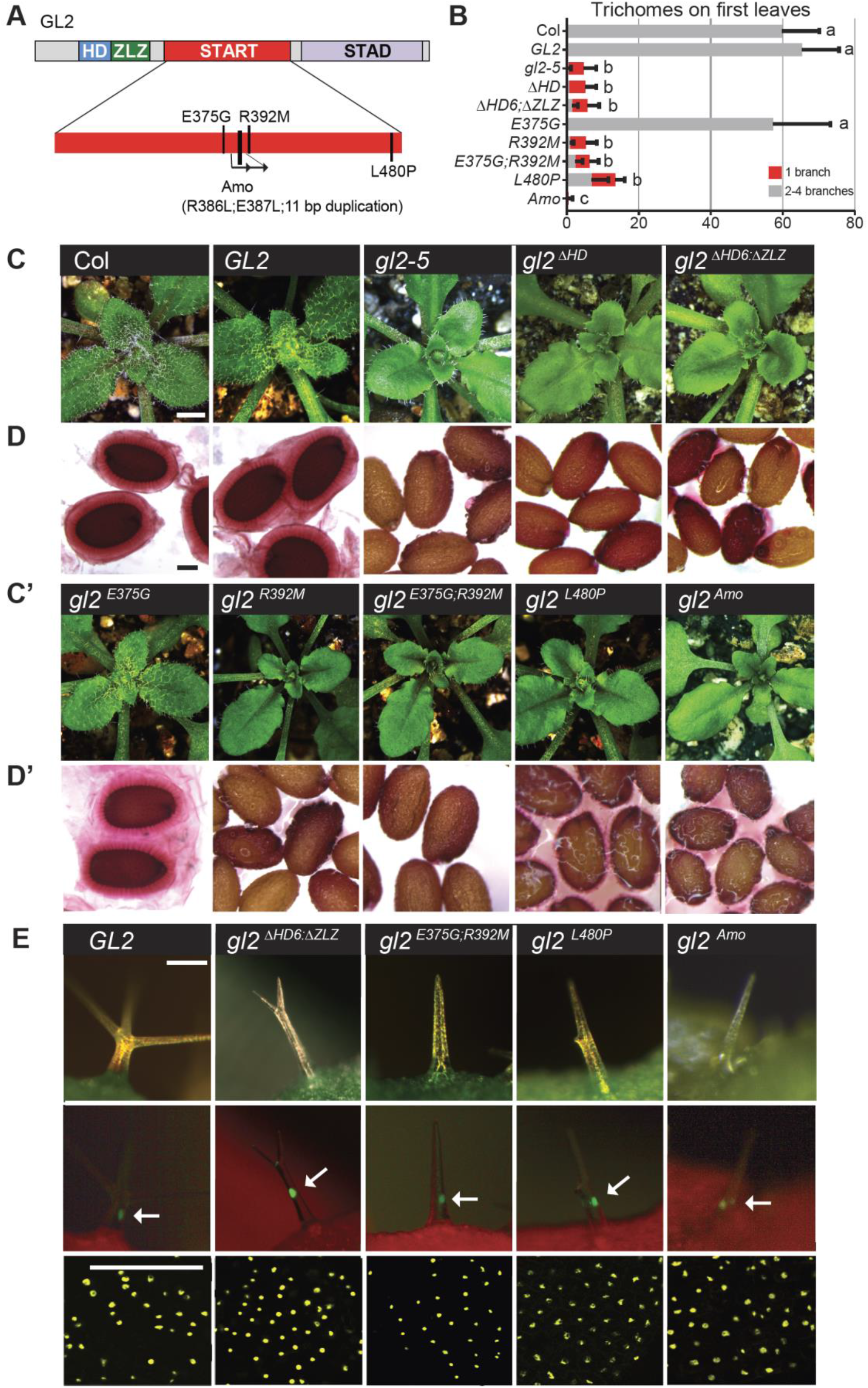
START domain missense mutants exhibit loss-of-function phenotypes despite nuclear localization of the corresponding EYFP-tagged proteins. **(A)** Schematic showing the functional domains of GL2. Positions of mutations within the START domain are indicated. See **Supplemental Figure 1B**. **(B)** Total number and branching pattern of trichomes on the first leaves. Gray bars indicate normal trichomes in plants expressing Col wild type, and EYFP-tagged GL2 and gl2^E375G^. Mutant *gl2-5* plants and those expressing EYFP-tagged gl2^ΔHD^, gl2^ΔHD6;ΔZLZ^, gl2^R392M^ gl2^E375G;R392M^, gl2^L480P^, and gl2^Amo^ exhibit unbranched trichomes (red bars). Values are mean ± SD for n = 20 plants. Letters denote significant differences from one-way ANOVA (p < 0.05) using Tukey’s multiple comparisons test. **(C)** Wild-type Col, *EYFP:GL2*, and *EYFP:gl2^E375G^* expressing rosettes display trichomes covering leaf surfaces. In contrast, *gl2-5* mutants as well as the EYFP-tagged HD mutants (*gl2^ΔHD^, gl2^ΔHD6;ΔZLZ^*) and START domain mutants (*gl2^R392M^, gl2^E375G;R392M^, gl2^L480P^*, and *gl2^Amo^*) exhibit trichome differentiation defects. Bar = 1 mm. **(D)** Ruthenium red staining of seeds to assay mucilage production. Wild-type Col, *EYFP:GL2*, and *EYFP:gl2^E375G^* seeds display a normal mucilage layer, while *gl2-5* mutants as well as the EYFP-tagged HD mutants (*gl2^ΔHD^, gl2^ΔHD6;ΔZLZ^*) and START mutants (*gl2^R392M^, gl2^E375G;R392M^, gl2^L480P^*, and *gl2^Amo^*) exhibit defective mucilage production. Bar = 100 μm. **(E)** Nuclear localization of EYFP-tagged GL2 and mutant gl2 transcription factors. Top panel: Imaging of leaf trichomes under white light (green, chlorophyll). Bar = 100 μm. Middle panel: Matching fluorescence images (red, chlorophyll) indicate nuclear localization of EYFP-tagged proteins in trichomes. Arrows mark the brightest nuclear signal in each image. Bar = 100 μm. Bottom panel: Confocal laser scanning microscopy of developing seeds reveals nuclear expression of EYFP-tagged GL2 and mutant proteins in seed coat cells. Bar = 50 μm.

We identified another missense allele, *W279R*, that maps to the N-terminus of the START domain, which also results in loss-of-function (**Supplemental Figure 3**). In *gl2^W279R^*, a highly conserved tryptophan is replaced with a charged residue within a predicted hydrophobic region of the binding pocket (**Supplemental Figure 1B and Supplemental Figure 3A**). In contrast, missense mutations in the predicted N-terminal α-helix of START (*gl2^A258P^, gl2^M271I^, gl2^I289N^, gl2^E294Q^*) do not significantly impair function (**Supplemental Figure 3A-E**)

Despite loss-of-function phenotypes, the four missense START alleles (*gl2^E375G;R392M^, gl2^Amo^, gl2^L480P^, gl2^W279R^*) and the *gl2*^Δ*HD6*;Δ*ZLZ*^ mutant retained nuclear localization of the correponding EYFP:tagged proteins (**Figure 3**, **Supplemental Figure 2B and Supplemental Figure 3F**). These results provide additional evidence that the HD and START domain are not required for nuclear location of GL2.

### HD, but not START domain, is required for DNA binding *in vitro*

HD-Zip IV TFs regulate the expression of various downstream target genes by binding to *cis*-elements in the respective promoter regions (Nakamura et al., 2006; Khosla et al., 2014). We performed electrophoretic mobility shift assays (EMSA) to examine whether DNA binding is regulated by the HD alone or by HD in combination with the START domain. For the DNA probe, we used the L1-Box element (TAAATGTA) implicated in GL2-DNA binding that is found upstream of several GL2 targets (Ohashi et al., 2003; Tominaga-Wada et al., 2009; Khosla et al., 2014). We produced recombinant Halo-tagged proteins using wheat germ *in vitro* transcription-translation. Despite repeated attempts, we were unable to obtain full-length recombinant GL2 protein that was functional in EMSA.

Given the high degree of sequence similarity between HD-Zip IV proteins (**Supplemental Figure 1**), we selected PDF2 and mutant versions analogous to our gl2 HD and START mutants (pdf2^K107E^, pdf2^K107T^, pdf2^ΔST^, pdf2^K107E;ΔST^, pdf2^K107T;ΔST^) for DNA binding studies. ATML1, which functions redundantly with PDF2, is reported to bind the octamer TAAATG(C/T)A (Abe et al., 2001). Thus, we performed EMSAs with a Cy3-labeled probe containing TAAATGTA (**Supplemental Table 3**). Both wild-type PDF2 and pdf2^ΔST^ caused a shift in probe mobility as an indication of DNA binding (**Figure 4A**). In contrast, the two missense mutations K107E and K107T resulted in defective mobility shift independently of the START domain. K107E completely abolished binding to the target sequence as indicated by a lack of mobility shift. While K107T resulted in partial DNA binding for both wild-type PDF2 (pdf2^K107T^) and pdf2^ΔST^ (pdf2^K107T;ΔST^) (**Figure 4A**). Western blotting confirmed uniform expression of the recombinant proteins used in our EMSA experiments (**Figure 4B**).

**Figure 4.**
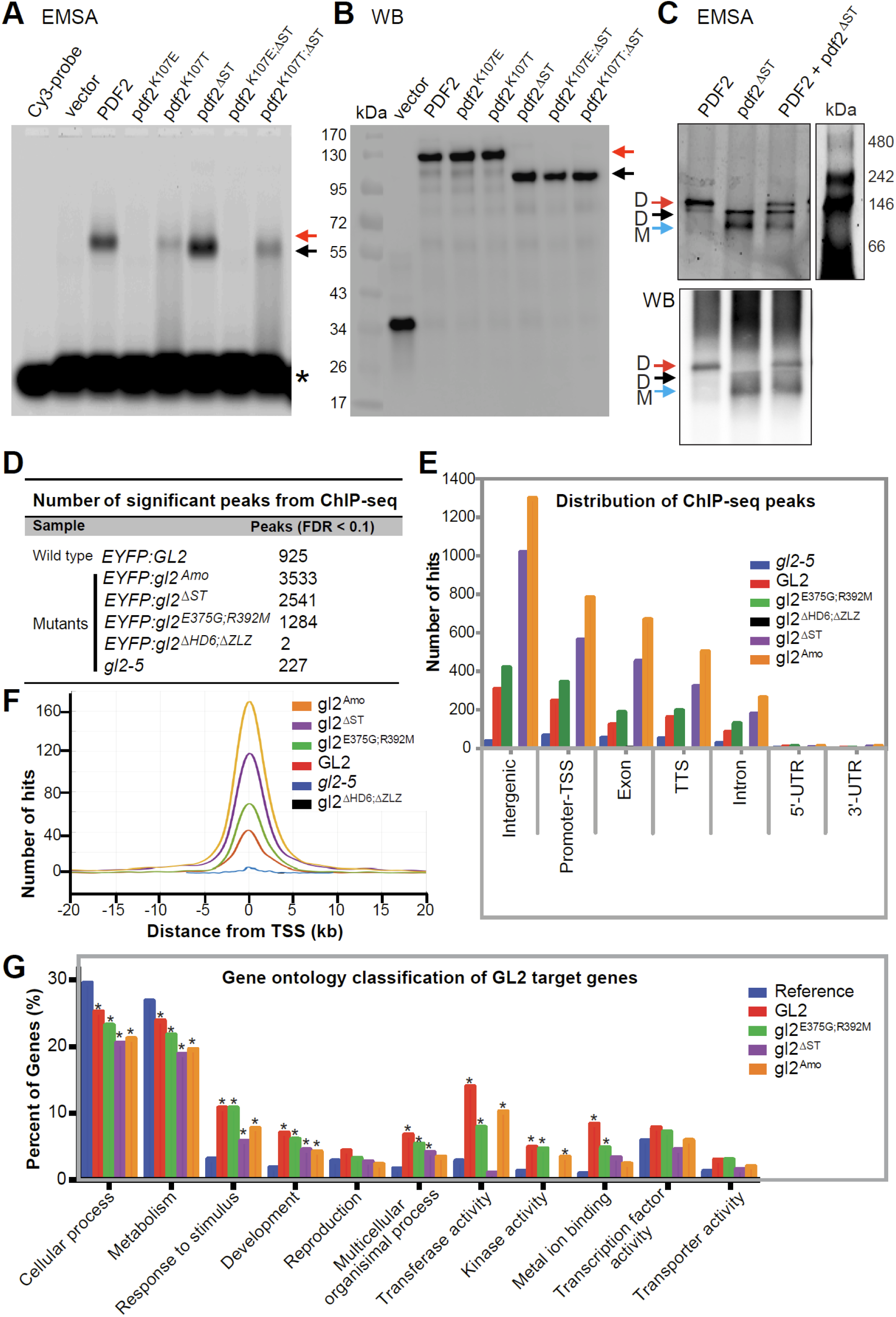
HD is required for DNA binding activity of PDF2 and GL2 while START domain is dispensable for DNA binding. **(A)** HD is required for *in vitro* DNA binding of PDF2 and pdf2^ΔST^ proteins. Electrophoretic mobility shift assays (EMSA) reveal band shifts for Halo:PDF2 (red arrow) and Halo:pdf2^ΔST^(black arrow), but not for proteins harboring HD mutation K107E. The K107T mutation results in partial DNA-protein complex formation indicated by a mixed population of bound and smeared molecules. Migration of the Cy3 probe containing the L1-box TAAATGTA is indicated (asterisk). Representative experiment from three independent experiments. **(B)** Western blot with anti-Halo Ab detects Halo-tagged proteins assayed for DNA binding in (**A**), representative of three independent experiments. **(C)** The pdf2^ΔST^ protein binds DNA as a monomer. EMSA coupled with polyacrylamide gel electrophoresis reveals that Halo:PDF2 binds the Cy3-labeled L1-box probe as a dimer (~230 kDa) (red arrow) while Halo:pdf2^ΔST^ binds DNA as a monomer (~90 kDa) (blue arrow) and weakly as a dimer (~180 kDa) (black arrow). Western blot (WB) (bottom) of the polyacrylamide gel with anti-Halo Ab detected Halo-tagged PDF2, and pdf2^ΔST^ at the expected migration positions matching that of the EMSA (top). Representative blot of two replicates. See **Supplemental Figure 4**. **(D-G)** Chip-seq data reveals that wild-type GL2 and mutants with compromised START domain function (gl2^ΔST^, gl2^Amo^, gl2^E375G;R392M^) are proficient in DNA binding *in vivo*. In contrast, DNA binding was abolished in the gl2^ΔHD6;ΔZLZ^ mutant. **(D)** Number of significant peaks (FDR < 0.1) from ChIP-seq data indicate that START domain mutants, but not the HD-ZLZ mutant, bind DNA target sites *in vivo*. **(E)** Genomic distribution of ChIP-seq binding events for wild-type GL2 and mutant versions are indicated (FDR < 0.1). TSS, transcription start site; TTS, transcription termination site; UTR, untranslated region. See **Supplemental Table 2**. **(F)** ChIP-Seq hit distribution from gene transcription start sites (TSS). The distribution of significant ChIP-seq peaks (FDR < 0.1) upstream and downstream from the TSS is shown for wild-type GL2 and mutants. See **Supplemental Figure 5**. **(G)** Functional classification of target genes from ChIP-seq data indicate over-represented GO terms from target genes whose promoters were bound by wild-type GL2 and mutants gl2^ΔST^, gl2^Amo^, and gl2^E375G;R392M^. Genes were classified based on molecular function and biological process, and a given gene may be represented in more than one category. Enrichment analysis was performed using AgriGO (Du et al., 2010). Asterisks indicate significant differences from the *Arabidopsis* genome reference (blue) (Fisher’s exact test, p < 0.01).

Combining EMSA with native polyacrylamide gel electrophoresis, we observed that the wild-type PDF2 protein binds the DNA as a dimer. In contrast, the pdf2^ΔST^ mutant binds the DNA target sequence as a monomer (**Figure 4C; Supplemental Figure 4**). The abnormal monomeric form was also seen in comparing the DNA-binding behavior of the mutant atml1^ΔST^ protein to wild-type ATML1 dimer (**Supplemental Figure 4**).

### START domain is dispensable for DNA binding *in vivo*

We performed ChIP-seq assays with seedlings expressing EYFP-tagged wild-type GL2 and mutant proteins to test whether the START domain is dispensable for DNA binding *in vivo*. The experiments included three START domain mutants (gl2^ΔST^, gl2^E375G;R392M^ gl2^Amo^) as well as the HD-ZLZ mutant (gl2^ΔHD6;ΔZLZ^), all of which display loss-of-function phenotypes similar to the *gl2-5* null mutant, but whose proteins are nuclear-localized (**Figures 1–3; Supplemental Figure 2**).

Data analysis revealed DNA binding proficiency for wild-type GL2 and all three START domain mutants whereas little or no binding was detected for gl2^ΔHD6;ΔZLZ^ and the *gl2-5* null mutant control (**Figure 4D; Supplemental Table 2**). Genomic distributions of binding sites were similar for wild-type GL2 and the START domain mutants. Most of the ChIP-seq hits were mapped to promoter or intergenic regions, consistent with the role of GL2 as a transcriptional regulator (**Figure 4E; Supplemental Table 2**). The relative positions of binding sites to transcription start sites (TSS) were similar for the wild-type GL2 and START domain mutants (**Figure 4F; Supplemental Figure 4**).

Gene ontology (GO) classification revealed overlapping functional classes of target genes bound by wild-type GL2 and the START domain mutants (**Figure 4G**). In the category “biological processes”, there was a similar enrichment for genes involved in cellular processes (GO:0009987; p = 4.8e-31) and metabolism (GO:0008152; p = 2.4e-42). In the category of “molecular function”, genes encoding transferase activity (GO:0016740; p = 8e-09), kinase activity (GO:0016301, p = 6.1e-07), and metal ion binding (GO:0046872; *p* = 4.5e-05) appeared as the most enriched genes for wild-type GL2 and two of the START mutants (gl2^Amo^, gl2^E375G;R392M^). Although the gene targets bound by the gl2^ΔST^ protein were similarly enriched for metal ion binding, they were not enriched for transferase activity and kinase activity (**Figure 4G**). Overall, the data indicate that START domain mutants retain DNA-binding activity in a manner that is comparable to wild type.

### START domain is required for GL2 homodimerization, while the HD is dispensible for this activity

Previous studies demonstrated the important role of the leucine zipper for homo- and hetero-dimerization of HD-Zip transcription factors (Landschulz et al., 1988; Sessa et al., 1993). Our EMSA experiments indicated that the START domain is critical for homodimerization and heterodimerization of PDF2 and ATML1 when bound to target DNA (**Figure 4C; Supplemental Figure 4**). We tested GL2 in yeast two-hybrid (Y2H) assays to further address the role of START in homodimerization of HD-Zip IV TFs. Unlike several other HD-Zip family members, GL2 does not show bait autoactivation in the Y2H assay (**Supplemental Figures 6A and 6C**). Y2H experiments confirmed that GL2 undergoes homodimerization but we did not detect heterodimerization of GL2 with other HD-Zip IV members nor was heterodimerization detected with two HD-Zip III members that were tested (**Supplemental Figure 6B**).

Next we assayed self interaction between different segments of GL2. Only the full-length GL2 protein and the segment encoding START+STAD displayed homodimerization. The START domain alone did not exhibit homodimerization, while the STAD domain alone could not be tested because of its bait autoactivation (**Figure 5A**). Consistent with the importance of the START domain for self interaction, deletion of START (*gl2*^Δ*ST*^) and two loss-of-function missense mutants (*gl2^R392M^* and *gl2^L480P^*) abolished homodimerization. The *gl2^E375G^* mutant, whose phenotype is similar to wild-type (**Figure 2**) supported dimerization (**Figure 5B**).

**Figure 5.**
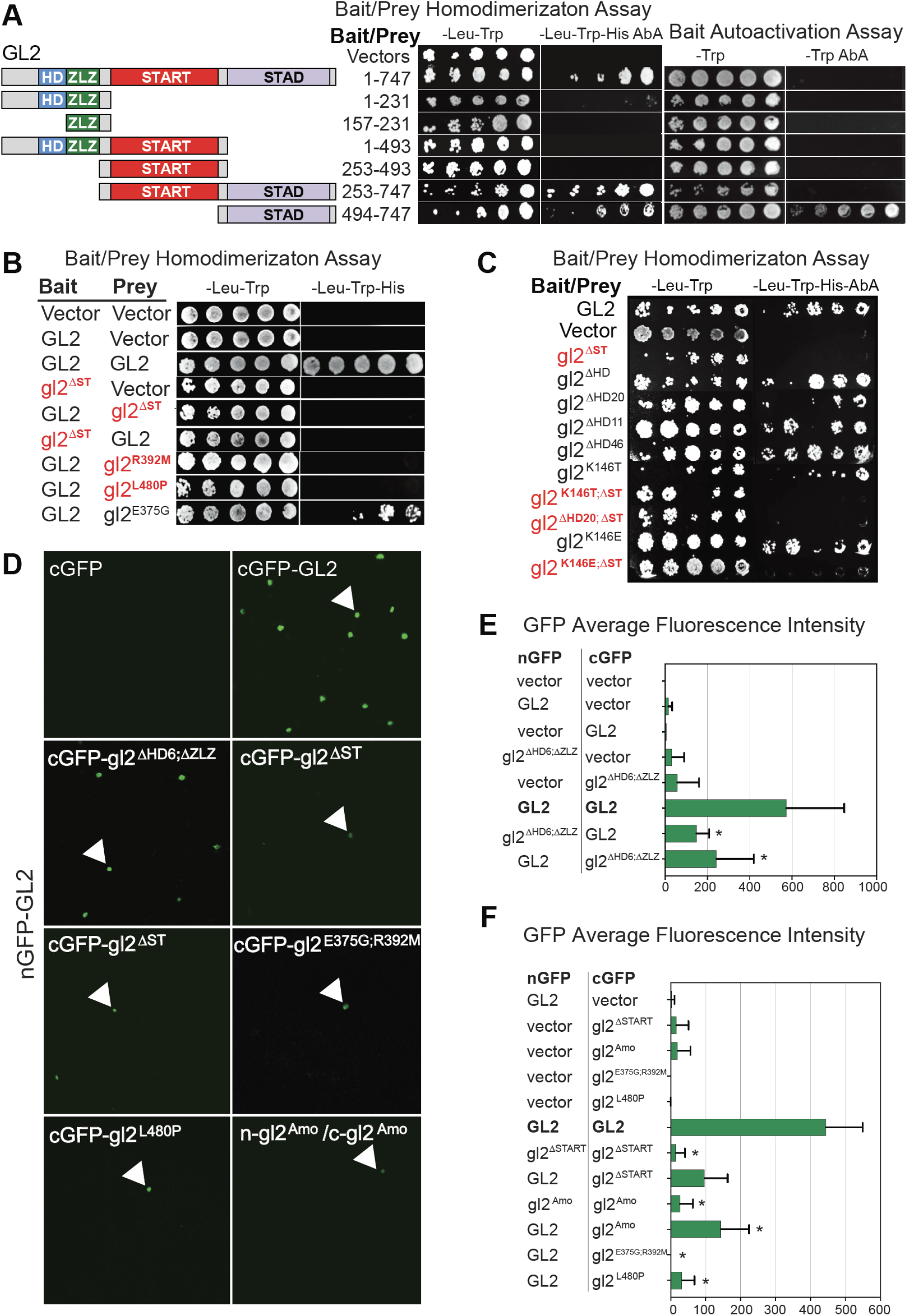
START domain is required for homodimerization of GL2. **(A)** Schematic representation of full-length GL2 and segments for Y2H assays (left). GL2 homodimerization of full-length protein (1-747) and START-STAD fragment (253-747) was observed (middle). The STAD fragment exhibits bait autoactivation (right). Homodimerization was assayed on -Leu-Trp-His Aureobasidin A (AbA) while bait autoactivation was assayed on -Trp AbA media. See **Supplemental Figure 7A**. **(B)** Deletion or missense mutations in the START domain disrupt homodimer formation in Y2H assays. GL2, gl2^ΔST^ and the indicated missense mutants were cloned into bait and prey vectors, co-expressed in yeast strain Y2HGold and assayed on the indicated growth media. Empty prey and bait vectors served as negative controls. **(C)** Y2H assays show that bait and prey fusion proteins interact to homodimerize only when START domain is present. Homodimerization was assayed on selective media as in **(A)**. HD mutants were proficient in homodimerization. START domain deletion resulted in loss of homodimerization in absence or presence of HD mutations (red). Wild-type GL2 bait and prey served as positive controls while empty bait and prey vectors represent negative controls. See **Supplemental Figures 6C-E and 7B**. **(D-F)** BiFC assays indicate homodimerization of wild-type GL2 but not HD-ZLZ mutant nor START domain mutants. **(D)** *N. benthamiana* leaf abaxial epidermal cells were transiently transformed with the indicated constructs and silencing suppressor p19. Interaction of full-length GL2 tagged with split GFP at the N-terminus (nGFP-GL2) with vector with the split GFP at C-terminus (cGFP), full-length GL2 and various gl2 mutants. Detection of gl2^Amo^ self-interaction is also shown. Nuclear expression is indicated by white arrows. Scale bar = 50 μm. See **Supplemental Figure 8**. **(E-F)** Quantitative analysis of BiFC assays in **(D)**. Mean pixel intensities from 10 images are shown. Error bars indicate standard deviations for two independent transformants in two trials. Asterisks indicate statistically significant differences (*, *p* < 0.0001 by *t* test).

We examined whether the HD plays a role in homodimerization. All of the HD mutants tested (*gl2^ΔHD^, gl2^ΔHD20^, gl2^ΔHD11^, gl2^ΔHD46^, gl2^K146T^, gl2^K146E^*) exhibited homodimerization but not bait autoactivation similar to wild-type *GL2* (**Figure 5C; Supplemental Figures 6C-E**). In contrast, double mutants with deletion of START (*gl2^K146T;ΔST^, gl2^ΔHD20;ΔST^, gl2^K146E;ΔST^*) abolished homodimerization (**Figure 5C**). These results show that GL2 dimerization is unaffected by HD mutations but is impaired by deletion of the START domain. Western blotting confirmed the expression of the various wild-type and mutant fusion proteins in yeast (**Supplemental Figure 7**).

Bimolecular fluorescence complementation (BiFC) assays in *N. benthamiana* (**Figure 5D-F**) verified our GL2 homodimerization results in yeast. Controls expressing either the N- or C-terminal half of GFP showed little or no fluorescence (**Supplemental Figure 8**). In contrast, strong fluorescent signals were observed in nuclei for samples expressing both halves of GFP fused to wild-type GL2, indicating homodimerization. Consistent with the reported role of the ZLZ domain in dimerization, the fluorescence intensity was significantly reduced when wild-type GL2 was tested for interaction with gl2^ΔHD6;ΔZLZ^ (**Figure 5E**). Reduced fluorescence intensity was similarly seen when wild-type GL2 was tested for interaction with four START domain mutant proteins (gl2^ΔST^, gl2^Amo^, gl2^E375G;R392M^, gl2^L480P^), as well as for self-dimerization for both of the START mutants we tested (gl2^ΔST^, gl2^Amo^) (**Figure 5F**). Taken together, the interaction assays in yeast and *N. benthamiana* demonstrate the critical importance of START for homodimerization of GL2, whereas the HD domain is not required for this activity.

### START domain, but not HD, regulates GL2 protein stability

Little is known about how the level of HD-Zip TFs is regulated post-translationally. It was reported that mutation of a putative lipid contact site in the START domain of ATML1 completely disrupts expression of the fluorescently tagged protein in lateral roots (Nagata et al., 2021). In our study, we observed that some EYFP-tagged gl2 proteins harboring mutations in either the HD or START, particularly gl2^W279R^, appeared weakly expressed in comparison to the wild-type protein (**Figure 3; Supplemental Figure 3F**).

To determine whether the HD and/or START domain play a role in regulating GL2 levels, we monitored protein half-life by treating *Arabidopsis* seedlings with cycloheximide, an inhibitor of protein synthesis. Proteins were extracted from seedlings at various intervals over a 10 or 24 h time course followed by immunodetection and quantification. We analyzed wild-type GL2, gl2^ΔHD20^ and gl2^ΔST^ along with three of the START missense mutants: gl2^W279R^, gl2^L480P^, and gl2^Amo^. As revealed by Western blot and protein quantification, wild-type GL2 and gl2^ΔHD20^ proteins remained stable over 24 h, whereas protein levels of various START domain mutants (gl2^ΔST^, gl2^L480P^, gl2^W279R^, gl2^Amo^) dropped rapidly (**Figures 6A and 6B; Supplemental Figure 9**). The gl2^ΔST^ and gl2^W279R^ mutant proteins were particularly unstable, having barely detectable protein levels at the 2-h time point (**Supplemental Figure 9**). In case of gl2^ΔST^, a degradation product of ~65 kDa also exhibited a diminished protein half-life (**Supplemental Figure 9**). We additionally examined the stability of proteins that harbored mutations in both the HD and START, but the low levels of expression made it difficult to interpret their half-lives in cycloheximide assays (**Supplemental Figure 10**). Taken together, our results provide strong evidence for the role of START in maintaining protein stability of GL2.

**Figure 6.**
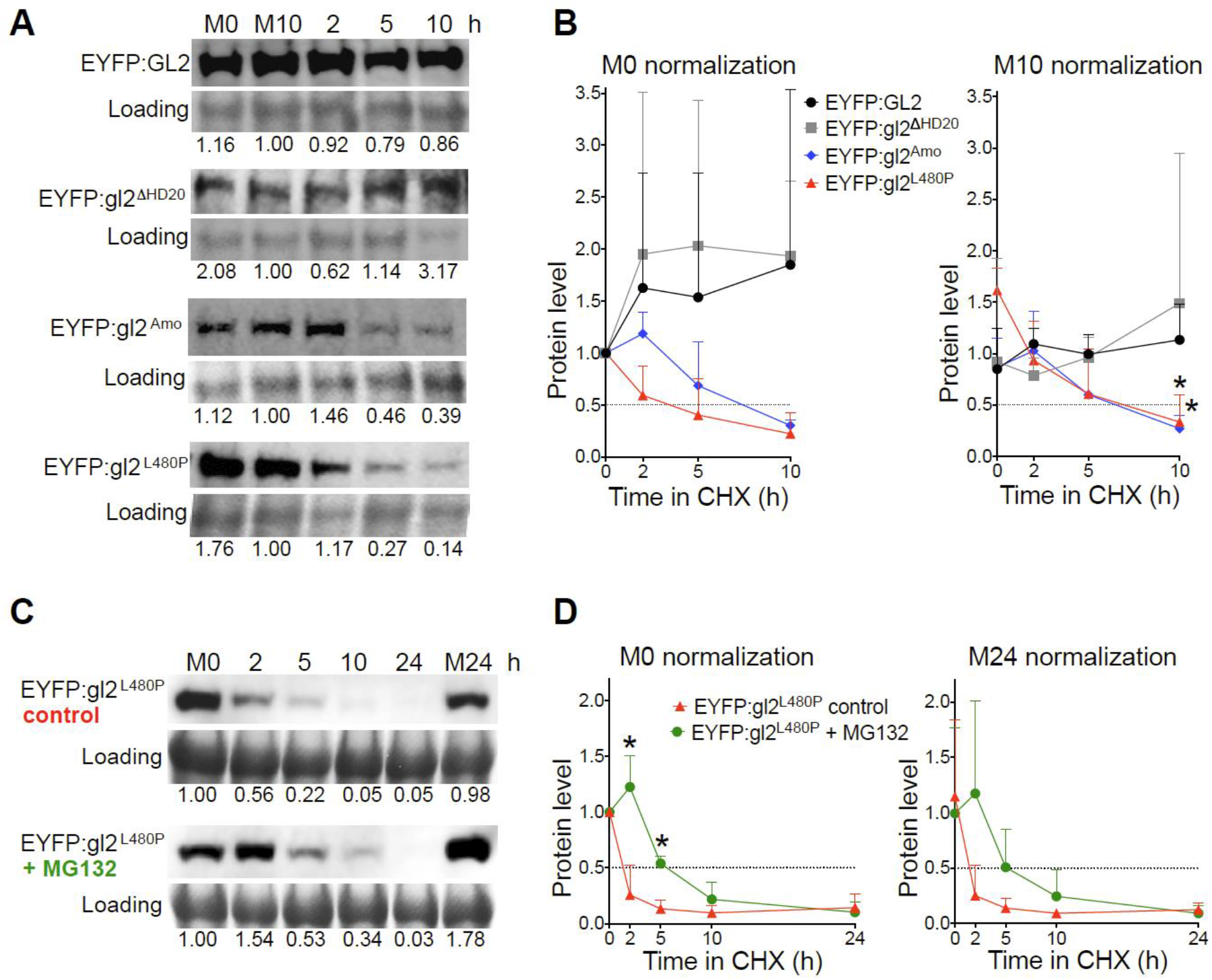
Cycloheximide experiments reveal that the START domain, but not HD, is critical for protein stability of GL2. **(A-B)** HD mutant gl2^ΔHD20^ behaves similarly to wild-type GL2, whereas START domain mutants gl2^Amo^ and gl2^L480P^ exhibit decreases in protein half-life. **(C-D)** Treatment with MG132 results in increased protein stability of START domain mutant gl2^L480P^. **(A and C)** Levels of EYFP-tagged proteins were monitored by Western blotting. Immunodetection with anti-GFP Ab was followed by Coomassie blue staining for loading controls. Mock treatments with DMSO at 0 (M0) and 10 (M10) or 24 (M24) h are indicated. Blots for each genotype are representative of three independent experiments. **(A)** Five-day-old seedlings expressing wild-type GL2 or mutant gl2 proteins were treated with cycloheximide (400 μM) for a 10 h time course. Values were normalized to loading controls and M10. See **Supplemental Figures 9 and 10**. **(B)** Five-day-old seedlings expressing EYFP-tagged gl2^L480P^ were treated simultaneously with cycloheximide (400 μM) and DMSO (control) or proteasome inhibitor MG132 (50 μM) for a 24 h time course. Values were normalized to loading controls and M0. **(B and D)** Protein quantification is shown for three Western blot experiments including that depicted in **(A)** and **(C)**, respectively. Values were normalized to M0 (left) or M10 (or M24) (right) controls. Protein half-life (0.5) is indicated by intersection with dotted lines. Error bars indicate SD for means from three experiments. Two-tailed *t*-test with unequal variance: *p ≤ 0.05.

To better understand the degradation pathway used to control protein levels, additional experiments were performed with one of the START mutants (L480P) in presence or absence of proteasome inhibitor MG132. Upon addition of MG132, the gl2^L480P^ protein remained stable for ~5-10 h during CHX treatment whereas the control treatment resulted in disappearance of the protein as early as 2 h (**Figures 6C and 6D**). The data suggest the possible involvement of Ubiquitin/26S proteasome system (UPS) in the regulation of GL2 protein levels.

## DISCUSSION

### Domain organization in HD-Zip IV TFs and nonoverlapping roles of HD and START

The combination of a DNA-binding HD with a lipid-sensing START domain in a single protein is plant-specific (Schrick et al., 2004) and likely represents a novel mode of action in a multidomain transcription factor. At the outset of this study, we envisioned that the START domain regulates transcription factor activity of HD-Zip IV TFs through its DNA binding domain. Our overall goal was to investigate the possible association or overlap in the roles of the HD and START domains. In addition to assaying phenotypic effects of genetic mutations *in planta*, we considered several aspects of transcription factor function, including subcellular localization, DNA binding, dimerization, and protein stability. We found that mutations affecting HD activity result in loss-of-function phenotypes, as illustrated by trichome differentiation defects, reduced seed mucilage and ectopic root hair formation similar to the null mutant *gl2-5, ΔST*, and multiple START missense mutants (**Figures 1 and 3; Supplemental Figure 3**). Both HD and the START domain are critical for epidermal differentiation, but the molecular mechanisms underlying their loss-of-function phenotypes are different. While mutations in the HD abolish DNA binding of the transcription factor, the loss of START domain function results in dimerization defects and unstable protein.

### Conserved role of HD in DNA binding

A missense mutation in the HD, changing a conserved lysine to a glutamic acid (K107E), completely disrupted DNA binding activity of PDF2 **(Figure 4A**). The analogous K-E substitution in the vertebrate HD transcription factors PITX2 and PITX3 also abolishes DNA binding (Saadi et al., 2001; Saadi et al., 2003; Sakazume et al., 2007). In contrast, we found that a K-T substitution at the same position (K107T) exerts a partial defect (**Figure 4A**). The effect of either of these two missense mutations on DNA binding was unaltered by the deletion of the START domain, consistent with our model that the HD functions in DNA binding independently of the START domain.

Our results indicate that START domain mutants are proficient in DNA binding under both *in vitro* and *in vivo* conditions (**Figure 4**). Several studies report that HD-Zip IV TFs bind DNA elements as dimers (Sessa et al., 1993; Palena et al., 1999; Tron et al., 2001). Surprisingly, our EMSA data suggested that START-deleted proteins bind DNA as monomers (**Figure 4C, Supplemental Figure 4**). While the START domain is not required for DNA binding, we confirmed the importance of the START domain for dimerization using self-interaction/homodimerization assays in yeast (**Figure 5B-C; Supplemental Figure 6**), and BiFC experiments *in planta* (**Figure 5D-F); Supplemental Figure 8**). Thus, the START domain behaves differently from the ZLZ domain, which was shown to be required for dimerization as well as DNA binding (Di Cristina et al., 1996).

### START domain is critical for protein stability

Our cycloheximide assays revealed that wild-type GL2 protein is relatively stable, while mutations in the START domain dramatically reduce protein half-life (**Figures 6A and 6B; Supplemental Figure 9**). It is noteworthy that gl2^L480P^, gl2^W279R^ and gl2^Amo^ harbor missense mutations in highly conserved amino acid residues predicted to be important for ligand binding (**Supplemental Figure 1B**). Thus, our findings support the hypothesis that ligand binding through the START domain stabilizes GL2 protein levels.

Stabilization of protein levels might aid in activation of the protein, promotion of protein–protein interactions, or correct protein folding. In eukaryotes, protein degradation acts as an important regulatory process that mediates optimum responses of cellular pathways to environmental cues. Our experiments with MG132 point to possible degradation of GL2 via the 26S proteasome (**Figures 6C and 6D**). However, proteasome inhibition is known to cause a global inhibition of translation resulting in proteome-wide effects (Larance et al., 2013). Thus, MG132 treatment might induce other pathways affecting protein stability, such as autophagy (Carlsson and Simonsen, 2015; Kang et al., 2018). Further investigation will be required to characterize the precise mechanism of GL2 protein degradation including possible post-translational modifications.

### Model for role of HD and START domain in promoting HD-Zip IV TF activity

We propose a model in which the function of the HD is to recruit the HD-Zip IV TF to the DNA target site (**Figure 7**). In this way, the multidomain transcription factor is correctly positioned on the DNA to activate or repress target genes. If HD function is missing, the HD-Zip IV protein is stably expressed and is capable of dimerization via its START domain. However, if the START domain is missing or non-functional due to mutation, the protein fails to dimerize and becomes unstable. Since START is required for both dimerization and protein stability, it is possible that these two functions are tightly interlinked. Thus far, we have not identified a mutant or condition in which these two processes are uncoupled. Therefore, START-domain-mediated dimerization likely contributes stabilization of the transcription factor. The sequence of events leading to transcriptional activity might involve several steps starting with ligand binding of the HD-Zip IV TF through START. Ligand binding could stabilize dimerization which in turn allows the transcription factor to bind the DNA via its HD. It is noteworthy that in the absence of the START domain, the monomeric form of the transcription factor binds to its DNA target site via its HD (**Figure 4C; Supplemental Figure 4**). In our model, unliganded monomers lacking the START domain are unstable, resulting in premature degradation.

**Figure 7.**
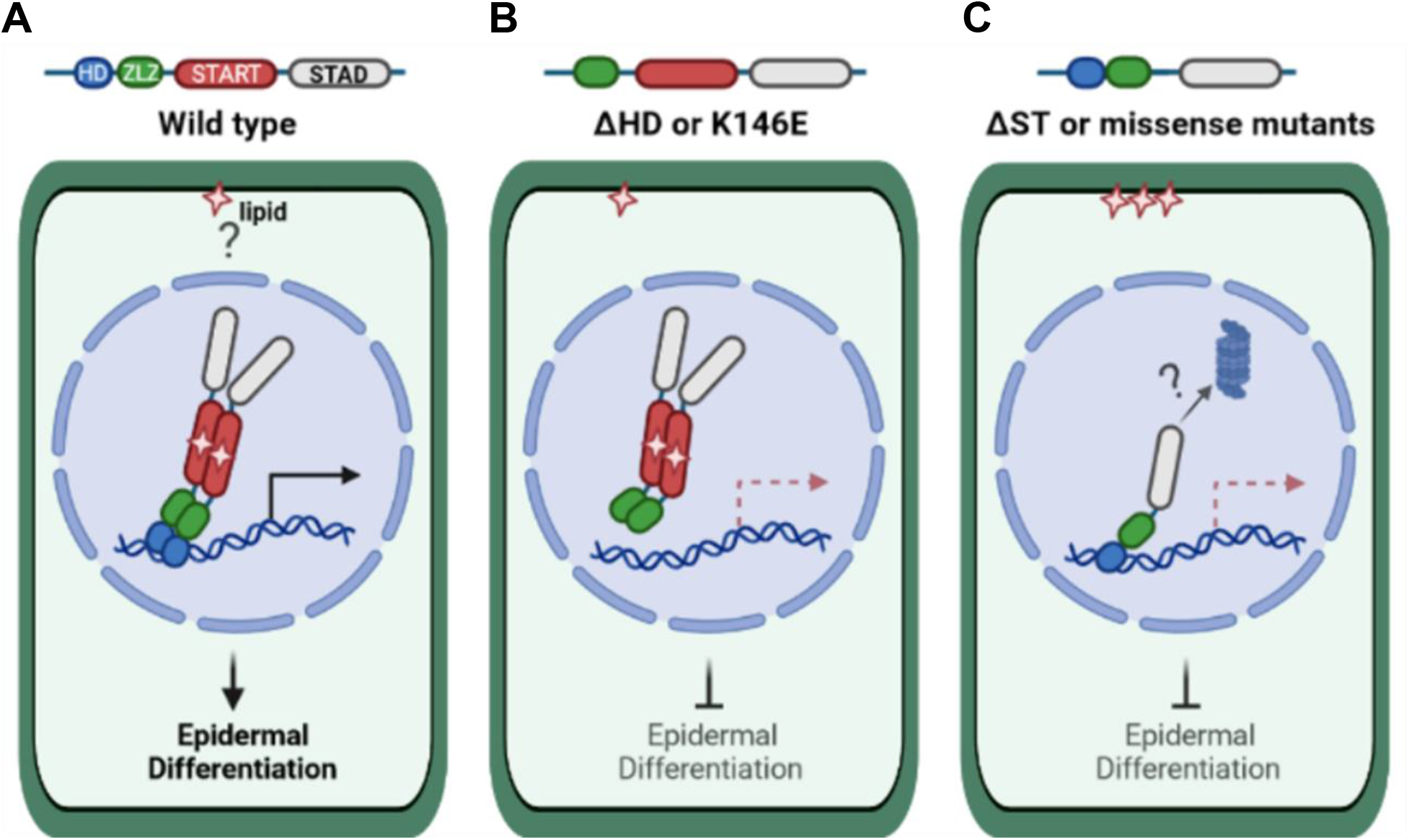
Model of HD and START domain roles in HD-Zip IV transcription factors. **(A)** Wild-type HD-Zip IV TFs, as exemplified by GL2, bind a lipid ligand that is possibly retrieved from a membrane compartment. The protein dimerizes and binds to DNA elements in the promoters of target genes, resulting in activation of transcription or repression of gene expression. For simplicity, only activation of gene expression is shown (horizontal arrow). Downstream target genes orchestrate cell differentiation of the plant epidermis. **(B)** Deletion or mutation of the HD (K146E in GL2) abolish DNA binding of the transcription factor, but dimerization and nuclear localization remain intact. Although the protein is found in the nucleus, epidermal defects in the corresponding mutants indicate that it is nonfunctional. **(C)** Deletion of or missense mutations in the START domain result in nonfunctional protein that is compromised in lipid binding. Nonetheless, the mutant protein is transported to the nucleus and binds DNA elements. Dimerization of the protein is affected, resulting in monomeric protein that is unstable, possibly due to degradation via a proteasome-dependent pathway.

### Other possible domain interactions in HD-Zip IV TFs

While our study was focused on the HD and the START domain, several other domains or motifs might also exert regulatory roles either independently or in conjunction with these functional domains. It is intriguing that while the START domain is important for homodimerization of GL2, it was unable to form dimers on its own and appears to require cooperative action with STAD (**Figure 5A**), a C-terminal domain of unknown function. Another candidate is the ZLZ domain, a plant-specific leucine zipper that contains ~10-20 amino acid loop with two highly conserved cysteines (Schrick et al., 2004). One study suggested that these cysteines are involved in redox control of dimerization as well as the regulation of DNA binding (Tron et al., 2002). Our results demonstrate that both the HD and START domain are dispensable for nuclear localization of the GL2 transcription factor in developing seeds and trichomes (**Figures 2 and 3**). An uncharacterized motif that maps elsewhere is likely responsible for nuclear localization of HD-Zip IV TFs. In previous studies, several plant HD-Zip I proteins were found to contain a nuclear localization signal (NLS) either at their amino- or carboxy-terminus (Arce et al., 2011; Cao et al., 2016; Shao et al., 2018).

Taken together, our study reports distinct roles of the HD and START domain, both of which contribute to the overall activity of HD-Zip IV TFs. These transcription factors are critical for orchestrating plant epidermal development, and they additionally play vital roles in responses to biotic and abiotic stress. Our results not only advance our current understanding of these developmentally important transcription factors but also lay the groundwork for engineering of plant-derived HD and START domains as modular units in synthetic biology applications. Future studies will explore how HD-Zip IV protein levels are tightly regulated during epidermal differentiation. Because these transcription factors also contain a specialized leucine zipper domain (ZLZ) and STAD, a plant-specific domain of unknown function, a promising avenue of future investigation will be to examine whether these domains work in association with HD and/or START to modulate various transcription factor activities.

## METHODS

### Plant material, growth conditions, and *Arabidopsis* transformation

Wild-type *Arabidopsis thaliana* ecotype Columbia (*Col*) and the *gl2* null allele *gl2-5* (Khosla et al., 2014) were used for this study. Seeds were stratified at 4°C for 3 days and grown on soil containing PRO-MIX PGX, vermiculite and perlite (Hummert International) in a ratio of 4:2:1 at 23°C under continuous light. The wild-type *proGL2:EYFP:GL2* and mutant *proGL2:EYFP:gl2* constructs were transformed into *gl2-5* plants using *Agrobacterium tumefaciens* GV3101 (pMP90) mediated floral dip (Clough and Bent, 1998). T1 progeny were screened on 20 μg/ml hygromycin B. Independent T1 transformants (21-55) were characterized for each genotype and 3:1 segregation for EYFP expression were confirmed in T2 progeny. Representative T3 homozygous lines with stable EYFP expression were confirmed by PCR genotyping and were used for further analysis.

### HD and START domain structural alignments

The closest structural homolog of the HD from GL2 was identified as the solution structure of *Gallus gallus* HOMEOBOX PROTEIN ENGRAILED-2 (Gg-En-2) (PDB ID = 3ZOB) using the SWISS-MODEL homology modeling server (Arnold et al., 2006; Benkert et al., 2011; Biasini et al., 2014). A structural homolog suitable for HD-Zip IV derived START domain modeling was identified as the crystal structure of human STARD5 (PDB ID = 2R55) using I-TASSER (Yang et al., 2015). Sequences from all 16 HD-Zip IV TFs from *Arabidopsis thaliana* (Nakamura et al., 2006) were aligned to corresponding structural homologs in Clustal O (Madeira et al., 2019). Consensus α-helices and β-sheets were visualized using ESPript 3.0 (Robert and Gouet, 2014).

### Plant constructs

The *GL2* cDNA was cloned into pENTR/D-TOPO using the pENTR/D-TOPO Cloning Kit (Invitrogen). Deletion and missense mutations *gl2*^Δ*ST*^, *gl2*^Δ*HD46*^, *gl2*^Δ*HD20*^, *gl2*^Δ*HD11*^, *gl2^K146T^*, *gl2^K146E^*, *gl2^A258P^*, *gl2^M271I^, gl2^W279R^, gl2^I289N^*, and *gl2^E294Q^* were generated using the Q5 Site–Directed Mutagenesis Kit (E0554S, New England Biolabs) following manufacturer’s protocol with primers listed in **Supplemental Table 3**. The *gl2*^Δ*ST*^ construct was used to generate *gl2*^Δ*HD20*;Δ*ST*^, *gl2*^*K146E*;Δ*ST*^ and *gl2*^*K146T*;Δ*ST*^ double mutants. After sequence confirmation, the wild-type and mutant pENTR/D-TOPO plasmids were transferred into the SR54 (*ProGL2:EYFP:GL2*) binary vector (Schrick et al., 2014). Inserts were amplified with gene specific primers using Q5 High Fidelity DNA polymerase (M0491, New England Biolabs), followed by purification of amplicons using the Nucleospin Gel and PCR Clean-up (Machery-Nagel) and *Dpn1* digestion. The vector and insert DNA fragments were assembled in a ratio of 1:5 using NEBuilder HiFi DNA Assembly (E5520S, New England Biolabs). The *gl2^L480P^, gl2^E375G^, gl2^R392M^*, *gl2^E375G;R392M^*, and *gl2^Amo^* mutants were generated by one-step PCR-based site-directed mutagenesis (Scott et al., 2002) using PfuUltra II Fusion HS DNA polymerase (Agilent Technologies). The Phusion^®^ Site-Directed Mutagenesis Kit (Thermofisher Scientific) was used to generate the *gl2*^Δ*HD*^ and *gl2*^Δ*HD6*;Δ*ZLZ*^ deletion mutants. Oligonucleotides for generation of mutants are listed in **Supplemental Table 3**. After verification by sequencing, the GL2 deletion constructs were restriction digested with *Kpn*I and *Sal*I and ligated into the binary vector SR54 (*proGL2:EYFP:GL2*) at the corresponding sites.

### Phenotypic assays and microscopy of *Arabidopsis*

*Arabidopsis* lines were assayed for leaf trichomes, root hairs and seed coat mucilage. Trichomes were quantified on first leaves from 20 individual plants. Leaf samples were placed on 0.8% agar-water plates and trichomes were counted manually upon viewing with a stereomicroscope. For root hair density analysis, seeds were vapor sterilized and sown on 0.5X MS media (Murashige and Skoog, 1962), followed by stratification at 4°C for 5 days. Seedlings were germinated in vertical orientation at 23°C under continuous light. Primary roots were imaged with a stereomicroscope 2-5 days after germination. For seed mucilage analysis, seeds were stained with ruthenium red (R2751, Sigma-Aldrich) following an established protocol (McFarlane, 2014) with slight modification. Briefly, 20-30 seeds per line were placed in a 24-well plate followed by addition of 800 μl 50 mM EDTA, and 2 h incubation with gentle shaking at room temperature. Seeds were washed to remove the EDTA and 800 μl 0.01% ruthenium red was added followed by 2 h incubation with gentle shaking. Seeds were washed with water (pH 7.5) and mounted on glass slides for imaging with a Leica DMIL LED compound microscope. Rosettes, roots, and trichomes were imaged with a Leica M165 FC stereomicroscope. Live imaging was performed on trichomes and developing seeds to detect EYFP expression using the stereomicroscope and Zeiss LSM 880 confocal microscope, respectively. Argon laser excitation of EYFP was at 488 nm, and the 515-550 band pass filter captured emission. To image EYFP-tagged proteins in primary roots, 4-d-old seedlings were stained with propidium iodide (10 μg/ml) to visualize cell boundaries. Seedlings were rinsed with distilled water and mounted in water under a coverslip. Images were collected using Zeiss LSM-5 Pascal confocal microscope. EYFP excitation/emission was at 488 nm/510-540 nm. Propidium iodide fluorescence was detected by excitation/emission of 543 nm/587-625 nm. Images were processed using Adobe Photoshop C4.

### *In vitro* transcription and translation

The full-length *PDF2* and *pdf2*^Δ*ST*^ cDNAs were transferred from pENTR/D-TOPO into destination vector pIX-HALO (CD3-1742, ABRC) using Gateway LR Clonase II Enzyme mix (ThermoFisher Scientific). The K107E and K107T mutations were generated using the Q5 Site-Directed Mutagenesis Kit (E0552S, New England Biolabs) with primers listed in **Supplemental Table 3** in pENTR/D-TOPO followed by transfer to pIX-HALO. The Halo fusion proteins were produced using 1.2 μg plasmid DNA of the respective pIX-HALO construct in a volume of 15 μl using the TNT SP6 High-Yield Wheat Germ Protein Expression System (L3260, Promega). Protein expression was confirmed by Western blot with Anti-Halo Tag monoclonal antibody (1:2000) (G9211, Promega).

### Electrophoretic mobility shift assay (EMSA)

Cy3-labeled and unlabelled dsDNA probes were generated with oligonucleotides listed in **Supplemental Table 3**. Annealing was performed with 25 μM oligonucleotides in 100 mM Tris-Cl (pH 7.5), 1M NaCl, and 10 mM EDTA at 95°C for 2 min, followed by 57°C for 5 min, 57-37°C transition over 90 min and 37°C for 2 min. EMSA reactions (20 μl) were prepared as previously described (Evens et al., 2017) with the following modifications: 6 μl of *in vitro* translated product was pre-incubated with binding buffer at 28°C for 10 min. Binding reactions were initiated by adding 200 nM of Cy3-labeled dsDNA probe, followed by a 20 min incubation. After electrophoresis at 4°C in a 0.6% agarose gel (1X TBE, pH 8.3) at 150 V for 1 h, or a 7.5% polyacrylamide gel (456-1025, Bio-Rad) at 100 V for 4 h, protein-DNA complexes were analyzed with a Typhoon™ Trio Imager (GE Healthcare) using the 580 nm emission filter for BP30Cy3, 600 PMT voltage, 532 nm green laser and high sensitivity.

### ChIP-seq assay

Seeds from EYFP-tagged *GL2*, *gl2*^Δ*ST*^, *gl2*^Δ*HD6*;Δ*ZLZ*^, *gl2^Amo^* and *gl2^R392M;E375G^* and *gl2-5* lines were surface sterilized and grown on 0.8% agar containing 1X MS (Murashige and Skoog, 1962) and 1% sucrose. Seeds were stratified for 3 to 5 d at 4°C and then transferred to 23°C and continuous light for 10 days. Seedlings were harvested and crosslinked with 1% formaldehyde for 10 min in a vacuum chamber. Glycine was added to a final concentration of 0.125 M, and the reaction was terminated by incubation for 5 min under vacuum. Chromatin was fragmented using a Covaris 220 focused ultrasonicator (Duty Cycle: 10%; Intensity: 5; Cycles per Burst: 200) to produce <500 bp fragments. ChIP experiment was performed as previously described (Khosla et al., 2014). Immunoprecipitated DNA was sequenced using Illumina sequencing technology. ChIP DNA was processed, prepared with 6-fold multiplexed Tru-Seq ChiP-seq libraries, and sequenced using short-read Illumina technology at the Kansas University Medical Center (KUMC) Genome Sequencing Facility. Libraries were sequenced (50 cycles) using the TruSeq Single Read Clustering Kit v3 and TruSeq SBS-HS v3 sequencing chemistry with index read.

### Bioinformatic analysis of ChIP-seq data

ChIP-sequencing was performed by Illumina HiSeq 2500 (Illumina, San Diego, CA) at a 50 base single read resolution. Read quality was assessed using FastQC (Andrews, 2010). Aligned read count statistics were obtained using the “flagstat” utility of the SAMtools suite (Li et al., 2009). Sequences were mapped to the *Arabidopsis thaliana* genome (TAIR10.24) using Bowtie2 (Langmead and Salzberg, 2012) (v 2.2.3; parameters: -q --phred33 --end-to-end --sensitive -p 16 -t). The mapping rates varied from a high of 97% to a low of 3%. Due to the low mapping rates of some replicates, the replicate aligned ChIP samples were merged (using “view” utility of the SAMtools suite (Li et al., 2009)) for greater coverage. The overall number of mapped reads for the merged ChIP samples varied from 20-70 million reads per sample (40 million reads per sample on average) except for the negative control sample, *gl2-5*, which did not contain express any EYFP-tagged protein, and had only 5 million mapped reads. See **Supplemental Data Set 1**. The merged ChIP samples were analyzed along with their corresponding input samples for significant peaks using MACS (Zhang et al., 2008) (v 1.4.2; parameters: -f BAM -g 0.135e9). Peaks calls passing a false discovery rate (FDR) cutoff of 0.1 were considered for downstream analysis.

### Target gene ontology analysis

AgriGO (http://bioinfo.cau.edu.cn/agriGO/index.php) was employed to extract functional annotations for genes whose promoter or intergenic regions were bound by the transcription factors (Du et al., 2010). Singular enrichment analysis (SEA) was applied to determine gene ontology (GO) term enrichment in each sample relative to the *Arabidopsis thaliana* TAIR9 reference. Analysis was performed using Fisher’s exact test (p < 0.01) with Bonferroni correction.

### Yeast two-hybrid (Y2H)

For **Figure 5A**, Bait and Prey constructs were prepared by restriction enzyme cloning with *Eco*RI and *Bam*HI inserts into the pGBKT7 and pGADT7 plasmids (Clontech). Bait constructs were transformed into the haploid yeast strain Y2HGold (*MATa trp1-901 leu2-3,112 ura3-52 hi3-200 gal4Δ gal80Δ LY2::GAL1_UAS_-Gal1_TATA_-Hi3 GAL2_UAS_-GAL2_TATA_-Ade2 URA3::MEL1_UAS_-Mel1_TATA_ AUR1-C MEL1*), and prey constructs were transformed into Y187 (*MATα ura3-52 hi3-200 ade2-101 trp1-901 leu2-3,112 gal4Δ gal80Δmet-URA3::GAL1_UAS_-Gal1_TATA_-LacZ MEL1*), using the LiAc method (Gietz and Woods, 2002), followed by selection on -Trp and -Leu, respectively. As a negative control, empty vectors were co-transformed with the desired constructs. Mating experiments were performed following Matchmaker Gold Yeast Two-Hybrid System (Clontech). Diploids were replica plated to media that selects for reporter gene activity of *HI3, AUR1-C*, and *MEL1*. To test for autoactivation of bait plasmids Y2HGold/ pDEST32 were plated in -Trp media and replica plated to media that selects for activity of the reporter genes *HI3* and *AUR1-C* respectively. All pair-wise combinations were investigated for protein-protein interaction by growing mated cultures to a density of OD_600_=1.0. Four serial dilutions at 4x each were prepared from the normalized cultures and spotted onto permissive and selective media for detection of interaction using a 48-pin multiplex plating tool (‘Frogger’, Dankar, Inc.). Plates were incubated at 30°C for 3-5 days, followed by imaging. Bait protein expression was confirmed by immunoblotting with Hela-c-Myc (9E10) polyclonal antibody (Gift from Stella Lee, 1:10 dilution).

For **Figure 5B-C and Supplemental Figures 6C-E**, full-length *GL2, PDF2, ATML1* cDNAs and the various deletion and missense mutations were transferred from pENTR/D-TOPO to pDEST32 bait and pDEST22 prey vectors (Proquest Two-Hybrid System, Invitrogen) using Gateway LR Clonase II Enzyme mix (ThermoFisher Scientific). For **Figure 5B**, both bait and prey constructs were transformed into Y2HGold, and interaction was assayed in this haploid strain. For **Figure 5B-C and Supplemental Figures 6C-E**, Y2HGold and the opposite mating type, Y187, were transformed with pDEST32 and pDEST22 constructs, using LiAc/PEG transformation, and plated on –Leu and -Trp media to select for the bait and prey plasmids, respectively. Mating experiments and selection for diploids were performed as described above. To test for autoactivation of bait plasmids Y2HGold/pDEST32 were plated in -Leu media and replica plated to media that selects for activity of the reporter genes *HI3* and *AUR1-C* respectively. Y2H assays for imaging were performed as described above. For analyzing bait protein expression, single colonies expressing pDEST32 constructs were grown at 30°C to saturation in media lacking leucine. Protein was extracted using the sodium hydroxide/trichloroacetic acid (NaOH/TCA) method and expression was monitored by Western Blot using a 1:500 dilution of GAL4 (DBD) RK5C1:sc 510-HRP mouse monoclonal antibody (sc-510HRP, Santa Cruz Biotechnology).

### Y2H quantitative β-galactosidase assay

For **Supplemental Figure 6A-B**, Gateway cloning was used to transfer genes for HD-Zip III and IV TFs from pENTR223.1 clones to pDEST22 and pDEST32 destination vectors (ProQuest Two-Hybrid System, Invitrogen). The bait and prey plasmids were transformed into yeast strain MaV203 (Invitrogen): *MATα leu2-3,112 trp1-901 his3Δ200; ade2-101 cyh2R can1R gla4Δ gal80Δ GAL1::lacZ HIS3_UASGAL1_::HIS3LEU2 SPAL10_UASGAL1_::URA3* following the Clontech Yeast Protocol Handbook. Four independent colonies from each transformation were selected to perform quantitative β-galactosidase liquid assay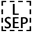 as previously described (Schrick et al., 2014).

### Bimolecular fluorescence complementation (BiFC) in *N. benthamiana*

To generate constructs for split GFP, Gateway compatible BiFC vectors 125-NXGW and 127-CXGW for N-terminal fusions of split GFP (a gift from Adrienne Roeder) were used for LR recombination reactions of the corresponding wild-type and mutant genes in pENTR/D-TOPO plasmids. Agroinoculation of *N. benthamiana* was performed as described in (Sparkes et al., 2006). Seeds of *N. benthamiana* were germinated on wet filter paper for 7d, and then the seedlings were planted on above described soil mix. For transient expression of fluorescent protein, leaves of 3–4-week-old plants grown at 22°C under 12 h light/12 h dark were used for agroinfiltration. Plasmids were electroporated into *A. tumefaciens* GV3101 and transformants were selected on agar plates containing the required antibiotics. Transformation of *N. benthamiana* leaves was performed with *Agrobacterium* taken directly from agar plates. Cultures were re-suspended in 10 mM MgCl2, 10 mM 2-(N-morpholino)ethanesulfonic acid pH 5.6 and 150 μM acetosyringone to an optical density of 0.2, and incubated for 2–4 h at 28°C in the dark. BiFC constructs were co-expressed with the p19 suppressor of gene silencing, derived from tomato bushy stunt virus (TBSV) (Voinnet et al., 2003). Leaves were infiltrated with a needle-less syringe into the abaxial side of leaves of 3–4-week-old *N. benthamiana* plants and examined after 2–3 d by confocal laser scanning microscopy using a Zeiss LSM-5 Pascal microscope configured for GFP excitation/emission of 488 nm/510-540 nm. Mean pixel intensities of three different circular regions of interest (ROIs) were quantified from 10 images for each construct combination using Image J software.

### *In vivo* protein stability assay

Wild-type and mutant seedlings were germinated on 0.5X MS media (Murashige and Skoog, 1962) for 4 days. On day 5, 30 seedlings of each genotype were transferred to 24 well-plates containing 1 ml 0.5X MS supplemented with 50 μM MG132 (474787, Sigma-Aldrich). Plates were sealed with surgical tape and placed on a shaker at 55 rpm for 16 h at 23°C under continuous light. After 16 h, the seedlings were washed 3X with 1 ml fresh media. Cycloheximide (C1988, Sigma-Aldrich) (400 μM in DMSO) or DMSO control was added at 0 h, and harvesting occurred at 0, 2, 4, 5, 8, 10 or 24 h, respectively. Samples of 30 seedlings were flash frozen in liquid nitrogen and stored at −80°C prior to protein extraction. Tissue was homogenized in liquid nitrogen, and hot SDS buffer (8 M urea, 2% SDS, 0.1 M DTT, 20% glycerol, 0.1 M Tris-Cl pH 6.8, 0.004% bromophenol blue) was added prior to SDS-PAGE and Western blotting. Anti-GFP (1:2000;11814460001, Roche) served as the primary Ab, followed by secondary Goat Anti-Mouse IgG [HRP] (1:3000; A00160, Genscript). Proteins were detected with SuperSignal West Femto Maximum Sensitivity Substrate (ThermoFisher Scientific) using an Azure 300 chemiluminescence imager (Azure Biosystems), and blots were stained with Bio-Safe Coomassie Blue G-250 (Bio-Rad) for loading controls. Band intensities were quantified with ImageJ software.

### Statistical Analysis

#### Trichome differentiation on the first leaves

Total number of trichomes and branching patterns were counted on the first leaves of 20 individual plants per genotype. Values are represented as mean ± SD. Ordinary one-way ANOVA using Tukey’s multiple comparisons test was applied to establish significant differences between genotypes (p < 0.05).

#### Root hairs of young seedlings

Numbers of root hairs on primary roots of 4-5 day-old seedlings were counted for 6-10 individuals per genotype. Values are represented as mean ± SD. Ordinary one-way ANOVA using Tukey’s multiple comparisons test was applied to establish significant differences (p < 0.05).

#### BiFC assays

For each combination of paired constructs, ten images were analyzed. Mean pixel intensity of three different circular regions of interest (ROIs) was quantified by the Image J software. An unpaired *t*-test was used to indicate statistically significant differences between wild-type and mutant using data from two independent experiments in two trials (*p* < 0.0001).

#### Protein stability assays

The graphed data represent two or three independent cycloheximide experiments with whole seedlings. Numerical values for protein levels were obtained from band intensity quantification and were normalized to the mock 10 (M10) and mock 24 (M24) samples depending on experiment, which were designated a value of 1.0. Protein half-life was determined by the protein level at a Y-axis value of 0.5. Values are represented as mean ± SD. *p < 0.05 indicates statistical significance using a two-tailed *t*-test with unequal variance.

### Accession Numbers

The ChIP-seq data is deposited in the NCBI Gene Expression Omnibus (Edgar et al., 2002) under the GEO Series accession number GSE186860 (https://www.ncbi.nlm.nih.gov/geo/query/acc.cgi?acc=GSExxx). The *Arabidopsis thaliana* genes described in this study are as follows: GL2 (At1g79840); PDF2 (At4g04890); ATML1 (At4g21750); ANL2 (At4g00730); FWA/HDG6 (At4g25530); HDG1 (At3g61150); HDG2 (At1g05230); HDG3 (At2g32370); HDG4 (At4g17710); HDG5 (At5g46880); HDG7 (At5g52170); HDG8 (At3g03260); HDG9 (At5g17320); HDG10 (At1g34650); HDG11 (At1g73360); HDG12 (At1g17920)

## Supporting information

Supplemental Data Set

## SUPPLEMENTAL INFORMATION

**Supplemental Figure 1**. Structural alignments of HD and START domain from HD-Zip IV transcription factors. Related to **Figures 1A and 3A**.

**Supplemental Figure 2**. Root hair phenotypes and nuclear localization in various START domain missense mutants. Related to **Figure 3**.

**Supplemental Figure 3**. Characterization of START domain missense mutants in the N-terminus of the START domain. Related to **Figure 3**.

**Supplemental Figure 4**. START domain is required for *in vitro* DNA binding as a dimer. Related to **Figure 4C**.

**Supplemental Figure 5**. ChIP-seq peak distance from transcription start site. Related to **Figure 4F**.

**Supplemental Figure 6**. START domain is required for GL2 homodimerization. Related to **Figure 5C**.

**Supplemental Figure 7**. Protein expression of Bait plasmids in Y2H experiments. Related to **Figure 5**.

**Supplemental Figure 8**. BiFC negative assays. Related to **Figures 5D-F**.

**Supplemental Figure 9**. Cycloheximide chase experiments reveal that the START domain is critical for protein stability of GL2. Related to **Figure 6**.

**Supplemental Figure 10**. Cycloheximide chase experiments with double mutants affecting both HD and START domain of GL2. Related to **Figure 6** and **Supplemental Figure 9**.

**Supplemental Table 1**. EYFP expression is independent of phenotypic rescue. Related to **Figures 2 and 3**.

**Supplemental Table 2**. Distribution and number of significant peaks from ChIP-seq data. Related to **Figure 4E**.

**Supplemental Table 3**. Oligonucleotides used in this study.

**Supplemental Data Set 1**. Mapping statistics of sequencing reads from ChIP-seq data.

## ACKNOWLEDGMENTS

This research was funded by the National Science Foundation (MCB1616818), National Institute of General Medical Sciences of the National Institute of Health under Award no. P20GM103418 (K-INBRE) and P2-RR-17686 (Confocal Microscopy Core, Kansas State University), USDA National Institute of Food and Agriculture Hatch/Multi-State project 1013013, and Johnson Cancer Research Center at Kansas State University. This is contribution no. 21-279-J from the Kansas Agricultural Experiment Station. We thank Joel Sannemann for help with confocal microscopy and Timothy Durrett for valuable scientific input.

## AUTHOR CONTRIBUTIONS

T.M., B.S., A.K. and K.S. designed the research, analyzed data, and wrote the paper. T.M. B.S., A.K., E.M.P and P.M.S performed experiments and analyzed data. T.M., R.L-R., A.L.W., E.M.P, A.K. and K.A.T. generated transgenic lines. S.G. performed ChIP-seq informatic analyses. K.S. oversaw the study.

