## Supplemental Figures and Tables for "START domain mediates *Arabidopsis* GLABRA2 transcription factor dimerization and turnover independently of homeodomain DNA binding"

**List of Supplemental Figures (10), Tables (3) and Data Set (1):**

Supplemental Figure 1
Supplemental Figure 2
Supplemental Figure 3
Supplemental Figure 4
Supplemental Figure 5
Supplemental Figure 6
Supplemental Figure 7
Supplemental Figure 8
Supplemental Figure 9
Supplemental Figure 10
Supplemental Table 1
Supplemental Table 2
Supplemental Table 3
Supplemental Data Set 1 (xls spreadsheet)

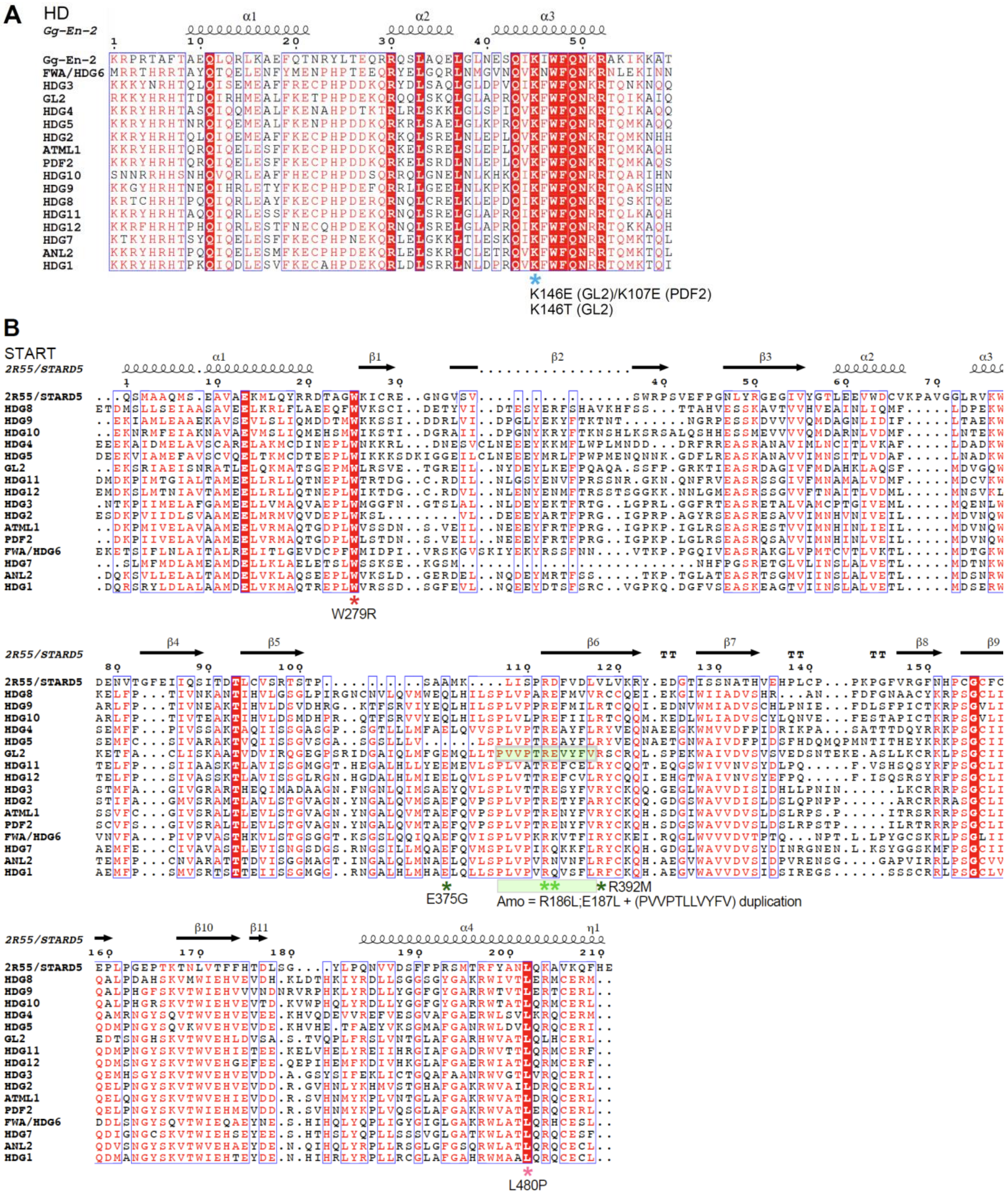
A
HD
Gg-En-2
α1
α2
α3
K146E (GL2)/K107E (PDF2)
K146T (GL2)
B
START
2R55/STARD5
W279R
E375G
R392M
Amo = R186L;E187L + (PVVPTLLVYFV) duplication
L480P
Gg-En-2
FWA/HDG6
HDG3
GL2
HDG4
HDG5
HDG2
ATML1
PDF2
HDG10
HDG9
HDG8
HDG11
HDG12
HDG7
ANL2
HDG1

**Supplemental Figure 1.** Structural alignments of HD and START domain from HD-Zip IV transcription factors. Related to **Figures 1A and 3A**.

**(A)** Structural alignment of homeodomain (HD) across all 16 members of the *Arabidopsis* HD-Zip IV transcription factor family with the solution structure for *Gallus gallus* HOMEOBOX PROTEIN ENGRAILED-2 (Gg-En-2) (PDB ID = 3ZOB) as reference. Consensus α-helices are displayed using ESPript 3.0 (Robert and Gouet, 2014). Similar amino acid residues are enclosed in blue boxes with identical residues highlighted in red, and numbering (1-59) for the HD only. Lysine targeted for missense mutations (K-E or K-T) is indicated (blue asterisk)

**(B)** Structural alignment indicates positions of START domain mutations examined in this study. Related to **Figure 3A**.

Structural alignment of the START domain across the HD-Zip IV family in *Arabidopsis* with the crystal structure for *Homo sapiens* STARD5 (PDB ID = 2R55) as reference. Consensus α-helices and β-sheets are displayed using ESPript 3.0 (Robert and Gouet, 2014). Similar amino acid residues are enclosed in blue boxes with identical residues highlighted in red. Numbering (1-211) corresponds to the START domain only. Positions of START mutations of GL2 analyzed in ChIP-seq (*E375G;R392M* and *Amo*) (**Figure 4D-G**) and the cycloheximide chase assays (W279R, Amo, and L480P) (**Figure 6 and Supplemental Figures 9 and 10**) are indicated below alignment (asterisks): *W279R* is a missense allele that adds a charged residue within a hydrophobic region of the binding pocket. *L480P* affects a lipid contact site in the C-terminal α-helix (Roostaee et al., 2009; Alpy and Tomasetto, 2014; Wojciechowska, 2021). In the *E375G;R392M* double mutant, a predicted salt bridge critical for ligand binding is disrupted (Roostaee et al., 2009). In *Amo*, the R386L and E387L missense mutations, which replace charged residues with leucines, are followed by an 11-residue duplication in a predicted ligand-contact region.

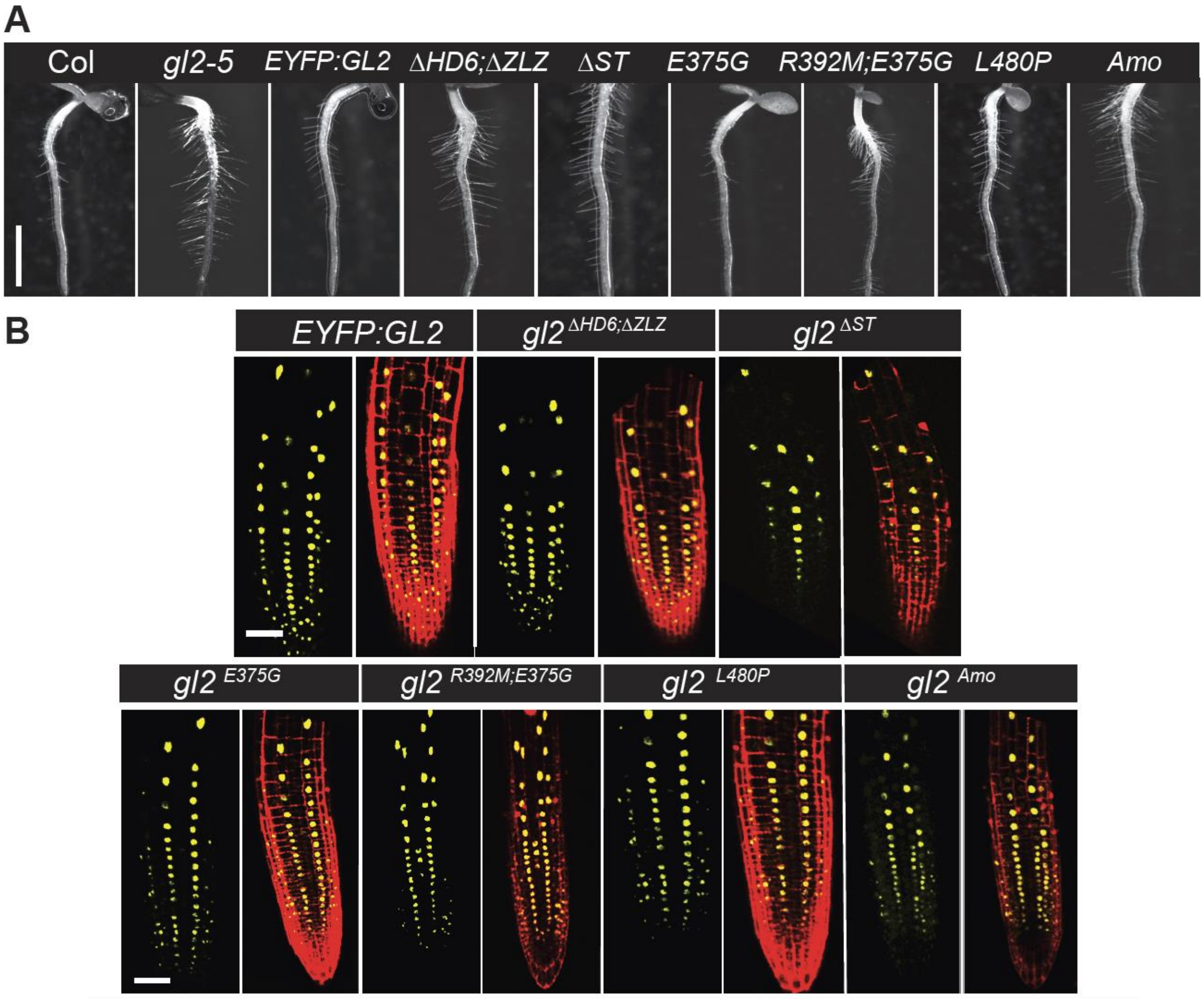


**Supplemental Figure 2.** Root hair phenotypes and nuclear localization in various START domain missense mutants. Related to Figure 3.

**(A)** Root hair phenotypes of 4-5 day-old seedlings. Bar = 1 mm. In contrast to wild-type Col, *gl2-5* null mutants display excess root hair formation. Expression of *EYFP:GL2* and *EYFP:gl2*$^{E375G}$ transgenes rescues the root hair phenotype of *gl2-5*. In contrast, aberrant root hair formation is observed for lines expressing EYFP-tagged gl2$^{\Delta HD6;\Delta ZLZ}$, gl2$^{\Delta ST}$, gl2$^{E375G;R392M}$, gl2$^{L480P}$ and gl2$^{Amo}$. Bar = 1 mm.

**(B)** Confocal laser scanning images of primary roots from 4-day-old seedlings expressing the EYFP-tagged proteins for lines depicted in **(A)**. Nuclear localization of the EYFP:GL2 wild-type and gl2 mutant proteins (yellow). Propidium iodide staining (red) was performed to visualize cell boundaries. Scale bar = 50 µm.

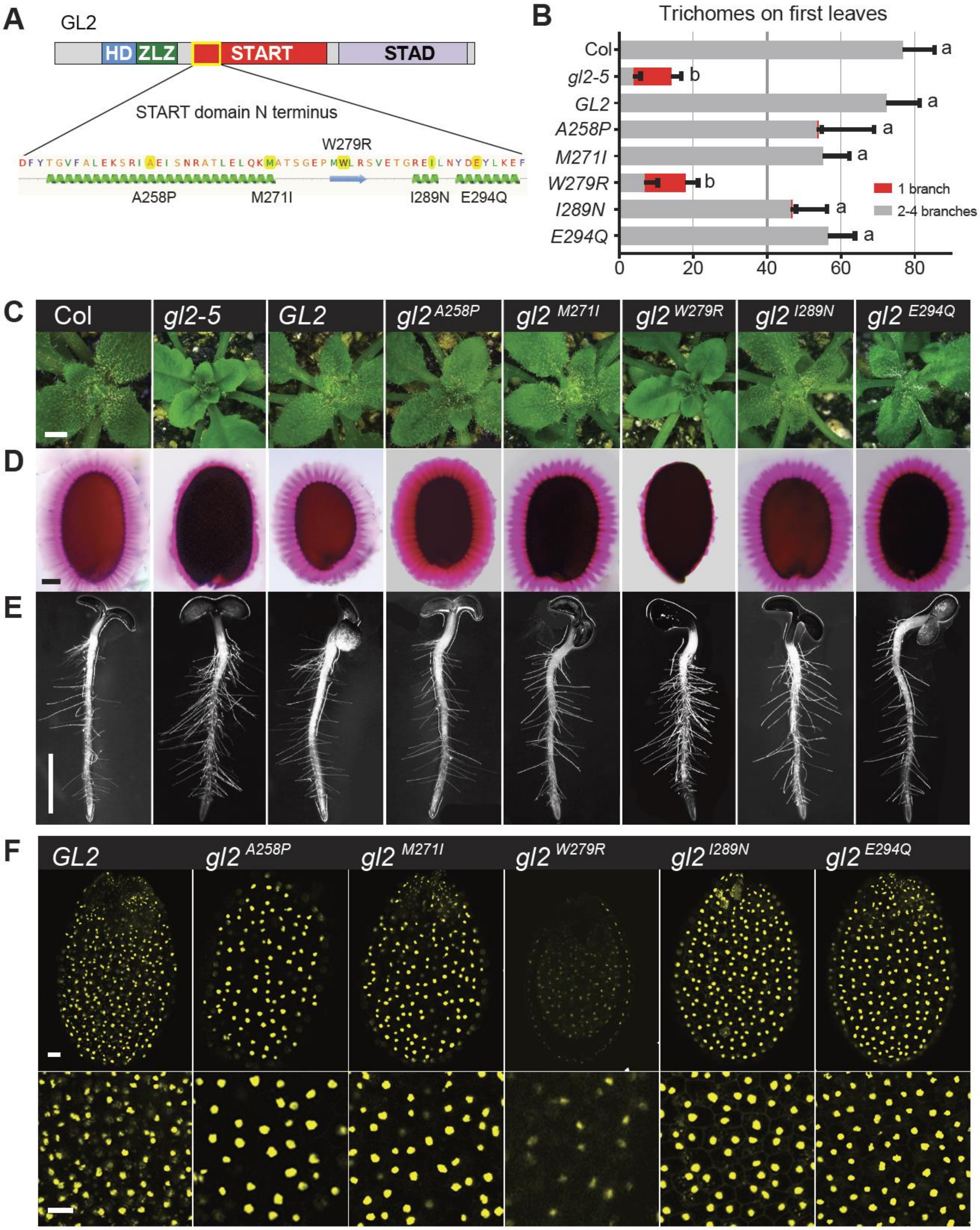
A
GL2
HD
ZLZ
START
STAD
START domain N terminus
W279R
A258P
M271I
I289N
E294Q
B
Trichomes on first leaves
Col
gl2-5
GL2
A258P
M271I
W279R
I289N
E294Q
1 branch
2-4 branches
0
20
40
60
80
C
D
E
F
Col
gl2-5
GL2
gl2 A258P
gl2 M271I
gl2 W279R
gl2 I289N
gl2 E294Q

**Supplemental Figure 3.** Characterization of START domain missense mutants in the N-terminus of the START domain. Related to **Figure 3**.

**(A)** Schematic showing the functional domains of the GL2 protein. Positions of mutations within the N-terminus of the START domain are indicated in relation to predicted helical (green) and strand (blue) secondary structures. See also **Supplemental Figure 1B** .

**(B)** Total number and branching pattern of trichomes on the first leaves. Gray bars indicate normal trichomes in plants expressing Col wild type, EYFP-tagged GL2 and the majority of the gl2 missense mutants shown in **(A)**. In contrast, mutant *gl2-5* plants and those expressing EYFP-tagged gl2$^{W279R}$ exhibit mostly unbranched trichomes (red bars). Values are mean ± SD for n = 20 plants. Letters denote significant differences from one-way ANOVA ($p < 0.05$) using Tukey's multiple comparisons test.

**(C)** Rosettes from wild-type Col, *EYFP:GL2*, and the majority of the gl2 missense mutants shown in **(A)** display trichomes covering leaf surfaces. The *gl2-5* mutants and the EYFP-tagged START domain mutant *gl2*$^{W279R}$ exhibit trichome differentiation defects. Bar = 1 mm.

**(D)** Ruthenium red staining of seeds to assay mucilage production. Wild-type Col, *EYFP:GL2*, and the majority of *EYFP:gl2* mutant seeds display a normal mucilage layer, while *gl2-5* mutants and the EYFP-tagged START domain mutant *gl2*$^{W279R}$ exhibit defective mucilage production. Bar = 100 μm.

**(E)** Root hair phenotypes of 4-5 d-old seedlings. In contrast to wild-type *GL2* and the majority of the *gl2* mutants, *gl2-5* and *gl2*$^{W279R}$ mutants exhibit excessive root hair formation. Bar = 1 mm.

**(F)** Nuclear localization of EYFP-tagged GL2 and mutant gl2 transcription factors. Top panel: Confocal laser scanning microscopy of developing seeds reveals nuclear expression of EYFP-tagged GL2 and mutant proteins in seed coat cells. Bar = 100 μm. Bottom panel: Magnified images. The EYFP:gl2$^{W279R}$ mutant protein exhibits weak expression in comparison to wild-type EYFP:GL2 and the other missense mutants. Bar = 50 μm.

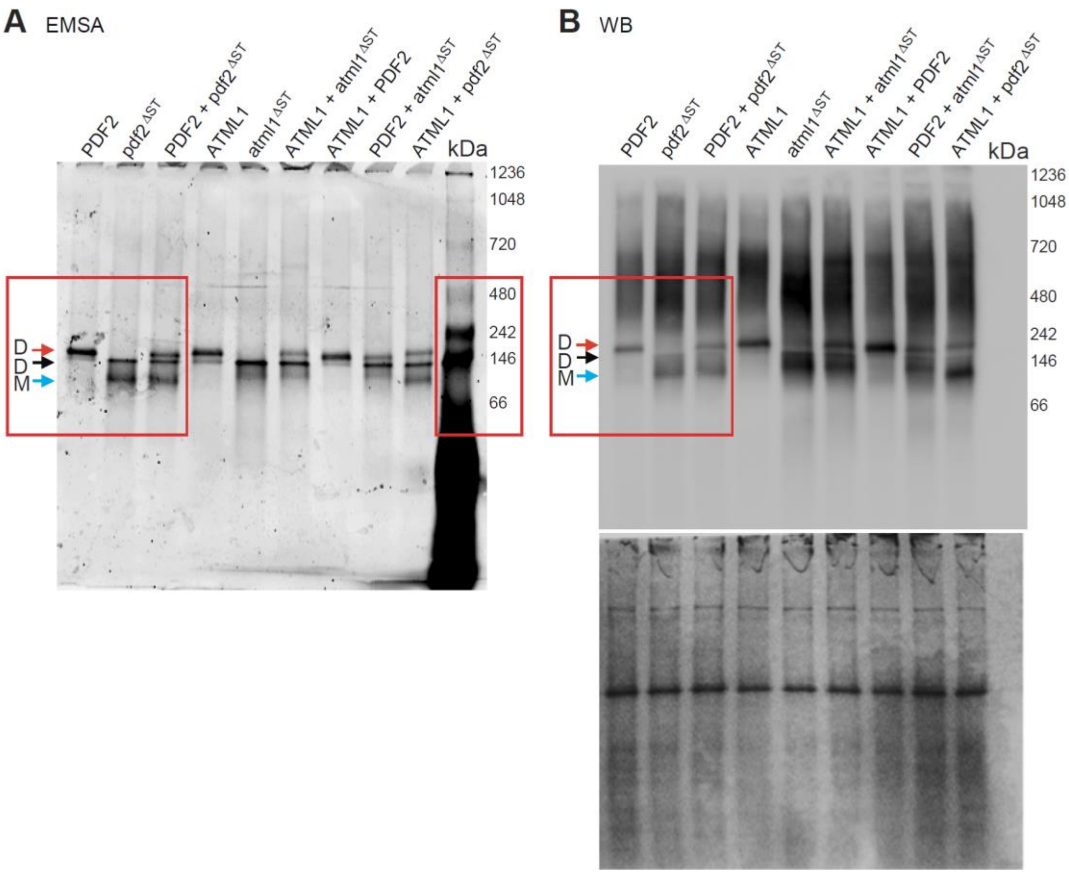


**Supplemental Figure 4.** START domain is required for *in vitro* DNA binding as a dimer. Related to Figure 4C.

**(A)** Electrophoretic mobility shift assay (EMSA) coupled with polyacrylamide gel electrophoresis reveals that Halo:PDF2 and Halo:ATML bind the Cy3-labeled L1-box probe as a dimer (~230 kDa) (red arrow) while Halo:pdf2$^{\Delta ST}$ and Halo:atml1$^{\Delta ST}$ bind DNA as a monomer (~90 kDa) (blue arrow) and weakly as a dimer (~180 kDa) (black arrow). A 1:1 mixture of Halo-tagged PDF2 and pdf2$^{\Delta ST}$, PDF2 and atml1$^{\Delta ST}$, as well as ATML1 and pdf2$^{\Delta ST}$, indicate additive banding patterns, suggesting that dimerization is affected when at least one of the two partners harbors a START domain deletion. Parts identified by red boxes are displayed in Figure 4C.

**(B)** Western blot (WB) (top) of the polyacrylamide gel in **(A)** with anti-Halo Ab detected Halo-tagged PDF2, ATML, pdf2$^{\Delta ST}$ and atml1$^{\Delta ST}$ at the expected migration positions as indicated from the EMSA **(A)**. After Western blotting, the blot was stained with Coomassie blue (bottom) to monitor equal loading of each EMSA reaction. The major band across all samples likely originates from wheat-germ extract used for *in vitro* transcription/translation of the Halo-tagged proteins.

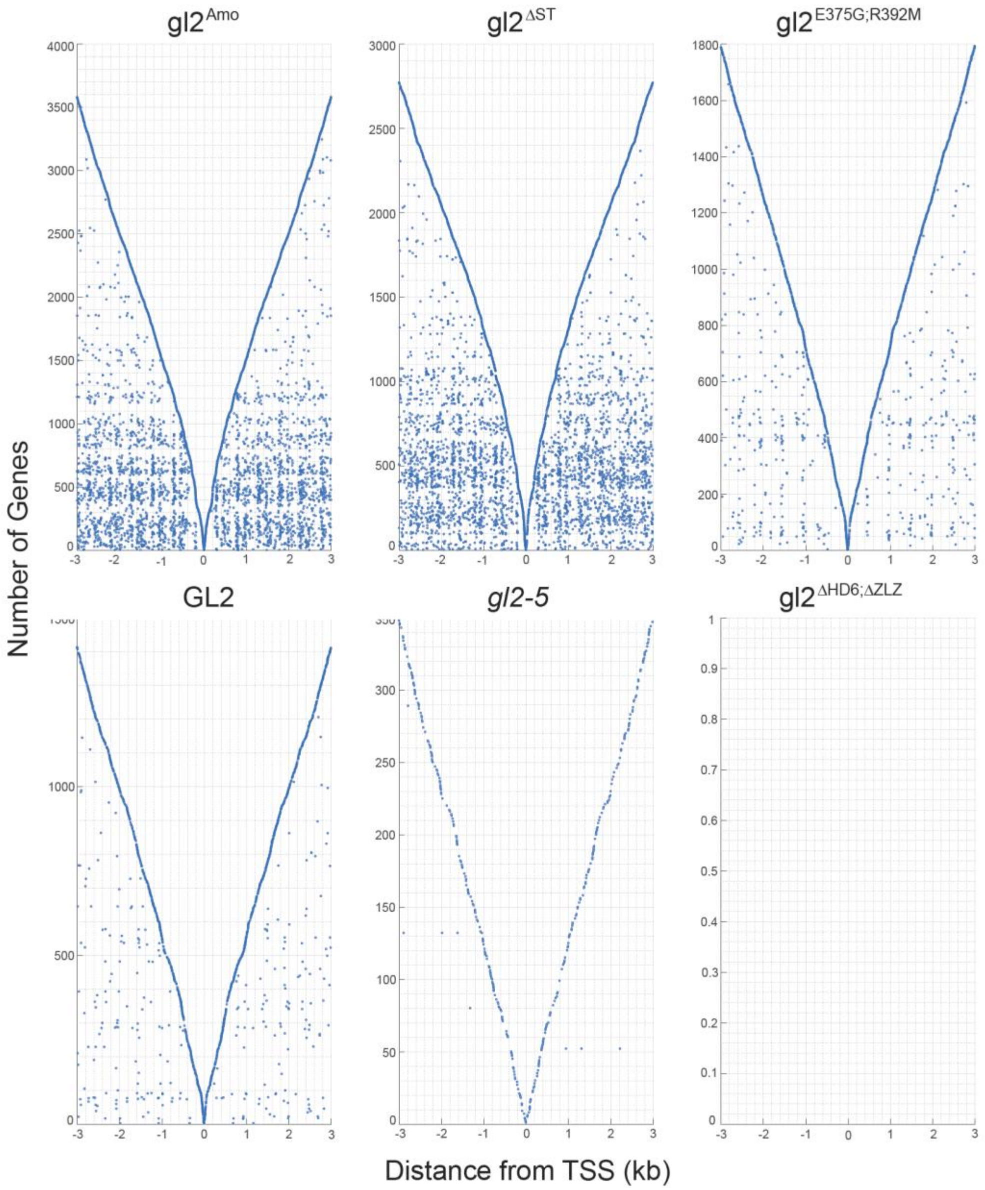


**Supplemental Figure 5.** ChIP-seq peak distance from transcription start site. Related to **Figure 4F**.

The significant ChIP-seq peaks (FDR < 0.1) from EYFP-tagged wild-type GL2 and mutant lines were plotted from gene transcription start sites (TSS). The genotypes are ordered according to the number of ChIP-seq peaks from left to right and top to bottom, with *gl2-5* representing the null mutant control that lacks EYFP. The Y axis represents individual genes ordered on the closest first hit to the furthest first hit in any of the samples within a 3 kilobase (kb) region from the TSS. Each dot represents a hit plotted for each gene along the X axis representing the distance from the TSS.

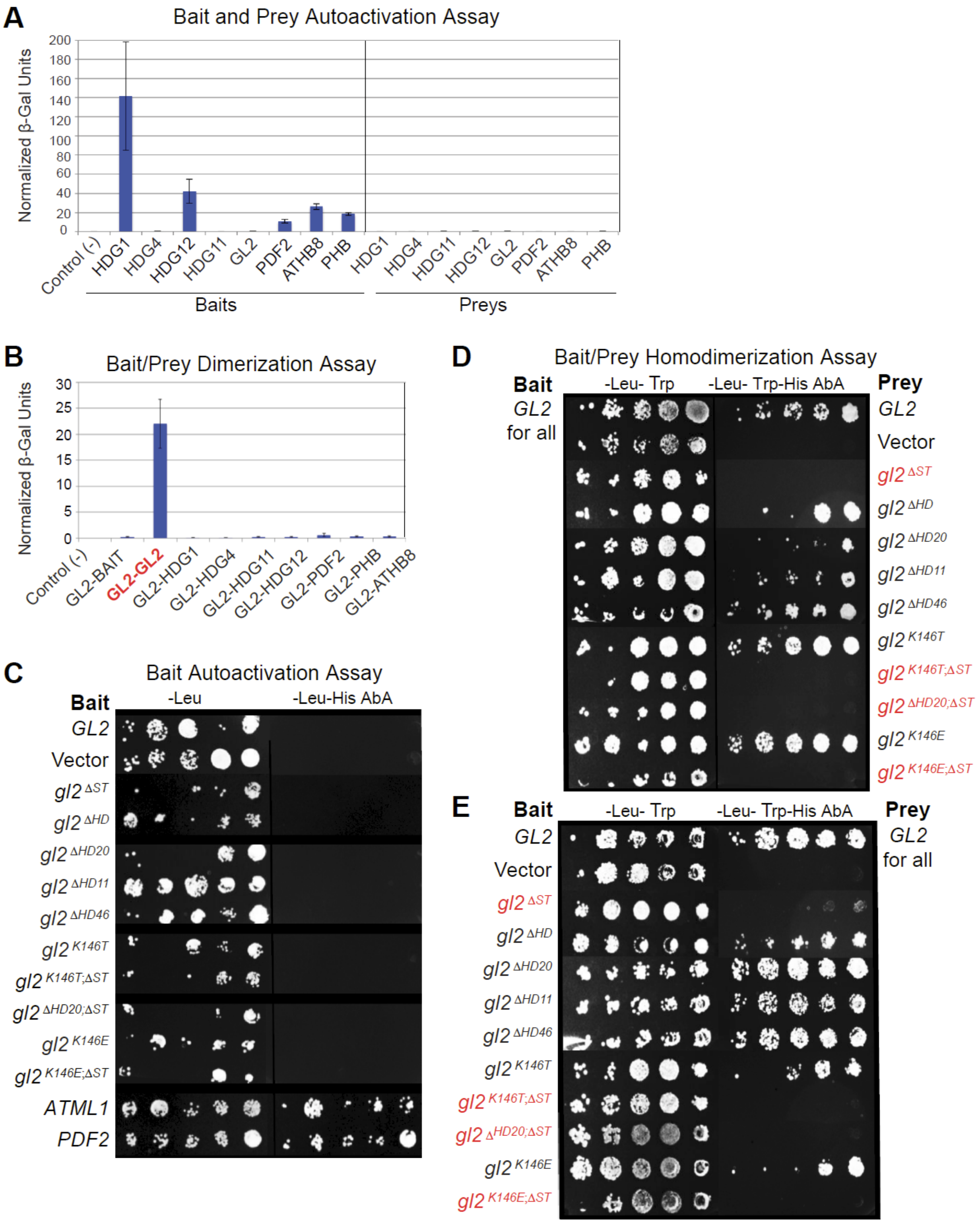


**Supplemental Figure 6.** START domain is required for GL2 homodimerization. Related to Figure 5C.

**(A and B)** Quantitative yeast ONPG assays for GL2 and other members of the HD-Zip TF family. Vector combinations used to co-transform yeast cells are given as prey/bait. Average β-galactosidase values were normalized to a positive control performed at the same time (pEXP$^{TM}$32/Krev1 and pEXP$^{TM}$22/RalGDS-wt, set to 100 β-units). The empty bait vector served as a negative control. Error bars indicate standard deviations for four independent transformants.

**(A)** GL2 does not display bait or prey autoactivation in Y2H assays. GL2, HDG4 and HDG12 do not exhibit bait autoactivation whereas all prey constructs tested do not exhibit autoactivation. Note: The HDG12 clone tested in this experiment was found to be truncated, lacking a portion of the START domain as well as C-terminal region.

**(B)** GL2 exhibits only homodimerization in Y2H assays. Heterodimerization was tested with other HD-Zip IV family members (HDG1, HDG4, HDG11, HDG12, PDF2) and HD-Zip III members (PHB, ATHB8).

**(C)** GL2 expression does not lead to bait autoactivation in contrast to expression of ATML1 or PDF2. GL2 wild-type and gl2 mutant bait constructs, as well as ATML1 or PDF2 bait constructs were transformed into haploid yeast cells. Autoactivation was monitored by growth on selective media (-Leu-His AbA). Lack of autoactivation was observed for wild-type GL2 and each of the mutants, while ATML1 and PDF2 exhibited autoactivation.

**(D)** Wild-type GL2 bait fusion protein homodimerizes with preys harboring HD mutations but not START domain deletion. The GL2 bait construct and indicated preys were transformed into yeast strains of opposite mating type.

**(E)** Wild-type GL2 prey fusion protein homodimerizes with baits harboring HD mutations but not START domain deletion. The indicated bait constructs and wild-type GL2 prey were transformed into yeast strains of opposite mating type.

**(D and E)** Haploid yeast were mated and diploids were grown on -Leu-Trp followed by selection on -Leu-Trp-His AbA to assay homodimerization. START domain deletion resulted in loss of homodimerization in absence or presence of HD mutations (red). Wild-type GL2 bait and prey served as positive controls, and empty bait and prey represent negative controls.

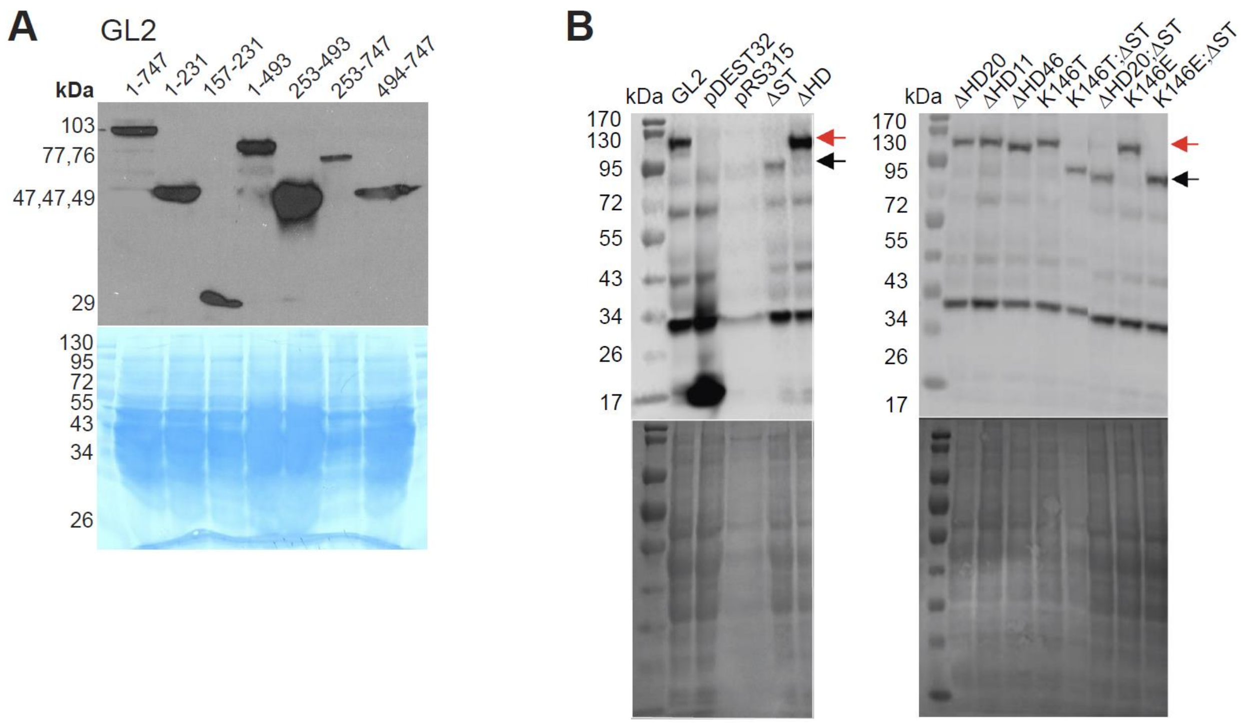


**Supplemental Figure 7.** Protein expression of Bait plasmids in Y2H experiments. Related to **Figure 5**.

**(A)** Western blotting confirms expression of bait proteins in **Figure 5A**. GL2 full-length protein and subdomains thereof are fused to the GAL4 DNA Binding Domain (DBD) and the human c-Myc epitope in the pGBK-T7 Bait vector. The primary antibody was c-Myc (9E10) (1:40) and the secondary antibody was Goat Anti-Mouse IgG [HRP] (A00160, GenScript) (1:3000). Bottom: Coomasie stained blot indicates loading of protein samples. Expected sizes of the GAL4 DBD-Myc fusion proteins are: 1-747, 103.3 kDa; 1-231, 47.3 kDa; 1-493, 76.6 kD; 253-493, 47.1 kDa; 494-747, 48.8 kD; 253-747, 75.7 kDa, 157-231, 29.4 kDa

**(B)** Western blotting with anti-GAL4(DBD) confirms expression of GL2 wild-type and mutant bait proteins in **Figure 5C**. Migration of wild-type and HD mutant proteins (red arrows) and START deleted proteins (black arrows) are indicated. pDEST32 is the vector control and pRS315 represents a second negative control. Coomassie blue staining after Western blotting indicates protein loading (bottom).

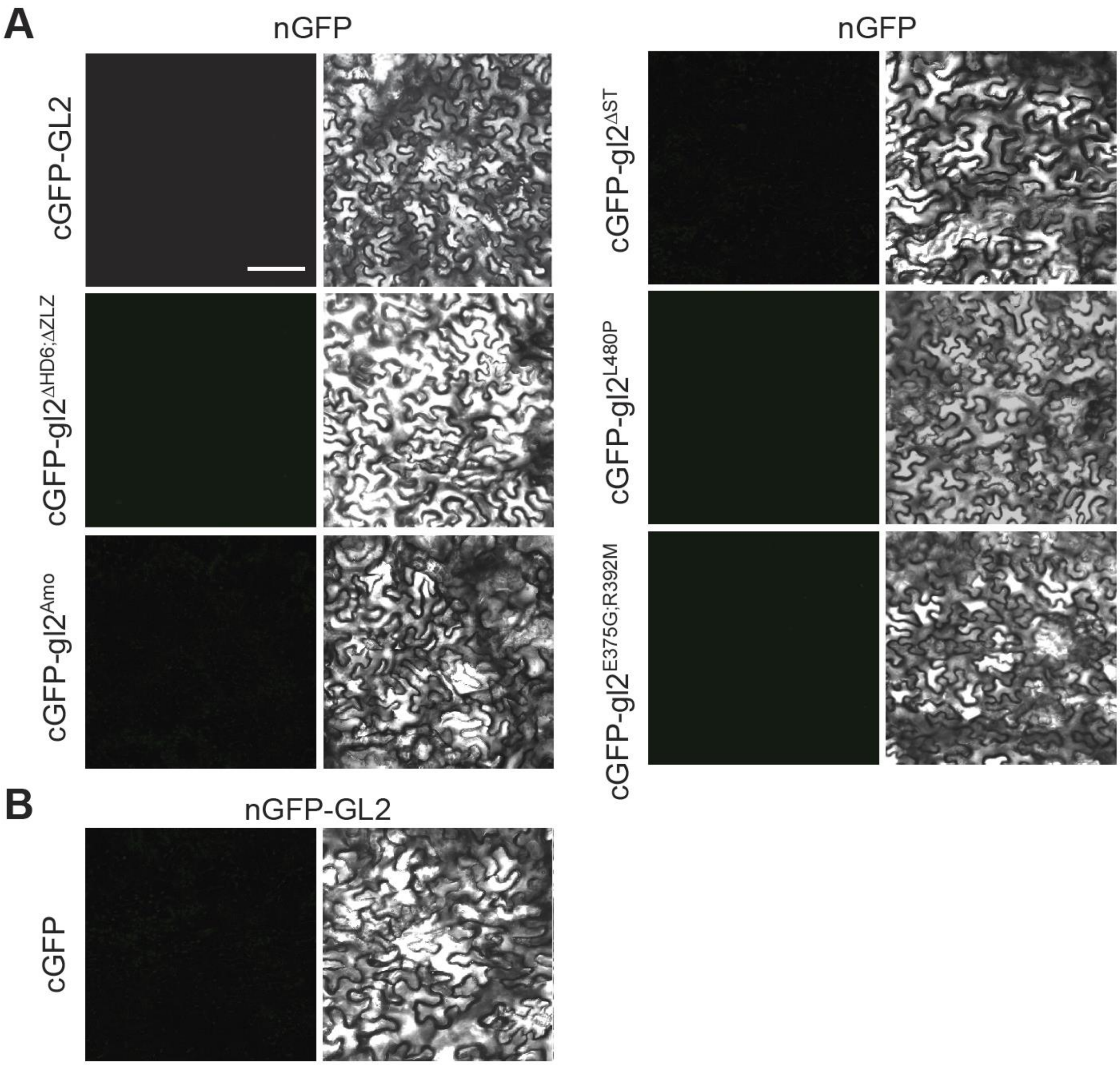


**Supplemental Figure 8.** BiFC negative assays. Related to Figure 5D-F.

(**A and B**) Confocal laser scanning microscopy of *N. benthamiana* leaves from BiFC experiments. Little or no fluorescence was observed in control experiments coexpressing the indicated proteins with empty vectors **(A)** nGFP (N-terminal half of GFP) and **(B)** cGFP (C-terminal half of GFP)

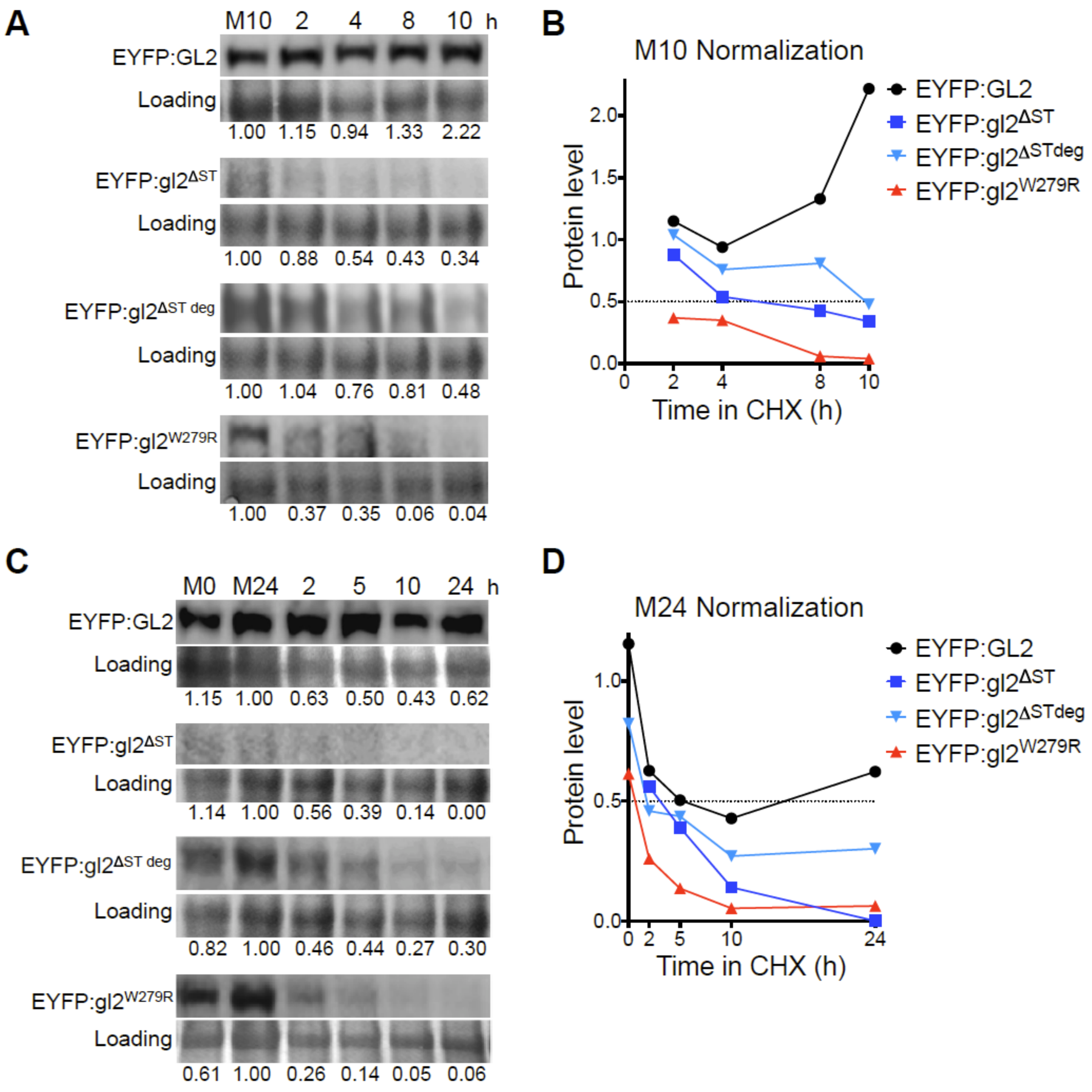


**Supplemental Figure 9.** Cycloheximide chase experiments reveal that the START domain is critical for protein stability of GL2. Related to **Figure 6.**

In comparison to wild-type GL2, START domain mutant protein $gl2^{\Delta ST}$ (and its degradation product $gl2^{\Delta ST\ deg}$) as well as $gl2^{W279R}$ exhibit a decrease in protein half-life.

Two independent experiments are shown in **(A-B)** and **(C-D)**, respectively. Five day-old seedlings expressing EYFP-tagged wild-type GL2 or START domain mutant gl2 proteins were treated with cycloheximide (400 μM) for a 10 h **(A-B)** or 24 h **(C-D)** time course.

**(A and C)** Levels of proteins were monitored by Western blotting. Immunodetection with anti-GFP Ab was followed by Coomassie blue staining for loading controls. Mock treatments with DMSO at 0 (M0), 10 (M10) or 24 (M24) h are indicated. The gl2$^{\Delta ST\ deg}$ protein was observed as a 65 kDa degradation product of gl2$^{\Delta ST}$.
**(B and D)** Graphs illustrate protein quantification for the experiments depicted in **(A)** and **(C)**, respectively. Values were normalized to loading controls and M10 **(A-B)** or M24 **(C-D)** controls. Protein half-life (0.5) is indicated by intersection with the dotted lines.

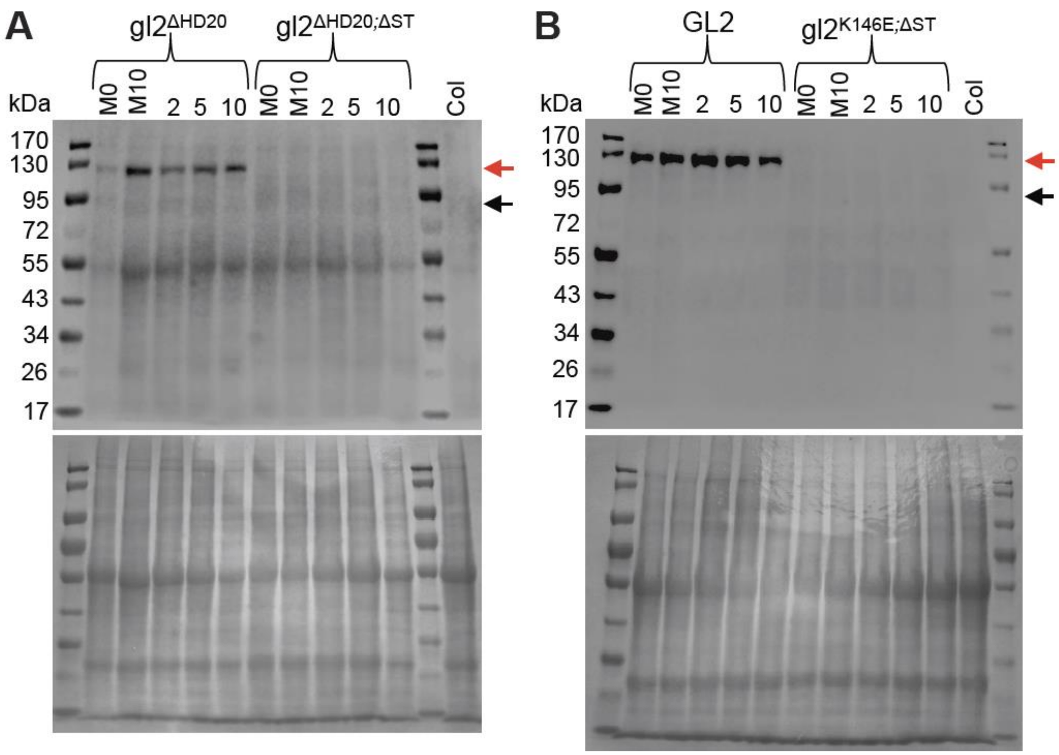


**Supplemental Figure 10.** Cycloheximide chase experiments with double mutants affecting both HD and START domain of GL2. Related to **Figure 6** and **Supplemental Figure 9**.

Five day-old seedlings expressing EYFP-tagged mutant gl2 or wild-type GL2 proteins were treated with cycloheximide (400 μM) for a 10 h time course. Mock treatments with DMSO at 0 (M0) or 10 (M10) h are indicated. Protein levels were monitored by Western blotting. Immunodetection with anti-GFP Ab (top) was followed by Coomassie blue staining for loading controls (bottom).

**(A)** The gl2$^{\Delta HD20}$ protein (expected size ~109 kDa; red arrow) is stable whereas the gl2$^{\Delta HD20;\Delta ST}$ protein (expected size ~83 kDa; black arrow) is barely visible or not detected over the 10 h time course.

**(B)** Wild-type GL2 protein (expected size ~111 kDa; red arrow) is stable while the gl2$^{K146E;\Delta ST}$ protein (expected size ~83 kDa; black arrow) is barely visible or not detected over the 10 h time course.

**Supplemental Table 1.** EYFP expression is independent of phenotypic rescue. Related to **Figures 2 and 3**.

Screening of T1 *gl2-5* transgenic plants for wild-type *EYFP:GL2* and mutant *EYFP:gl2* constructs indicated no simple correlation between the trichome rescue phenotype and the intensity of EYFP expression in the nucleus. All wild-type *EYFP:GL2* lines exhibited phenotypic rescue although 5 out of 28 lines exhibited low levels of EYFP expression.  Both HD mutants and START deletion mutants failed to rescue but showed varying levels of EYFP expression. The gl2$^{K146T}$ mutants displayed a partial phenotype. n = 28-50 T1 plants for each of the constructs.

| **Genotype** | **n** | **Level of phenotypic rescue** | | | **EYFP nuclear expression** | | | |
|---|---|---|---|---|---|---|---|---|
| | | Complete | Partial | None | High | Medium | Low | Undetected |
| *GL2* | 28 | 27 | 1 | - | 22 | 1 | 5 | |
| *gl2*$^{\Delta ST}$ | 50 | | | 50 | | | 30 | 20 |
| *gl2*$^{\Delta HD46}$ | 37 | | | 37 | 37 | | | |
| *gl2*$^{\Delta HD20}$ | 30 | | | 30 | 11 | 14 | 5 | |
| *gl2*$^{\Delta HD11}$ | 34 | | | 34 | 11 | 22 | 1 | |
| *gl2*$^{K146T}$ | 35 | 19 | 7 | 9 | | 32 | 3 | |
| *gl2*$^{K146E}$ | 55 | | | 55 | | 41 | 13 | 1 |
| *gl2*$^{K146T:\Delta ST}$ | 47 | | | 47 | | | 32 | 15 |
| *gl2*$^{K146E;\Delta ST}$ | 46 | | | 46 | | | 24 | 22 |
| *gl2*$^{\Delta HD20;\Delta ST}$ | 55 | | | 55 | | | 28 | 27 |

**Supplemental Table 2.** Distribution and number of significant peaks from ChIP-seq data. Related to **Figure 4E**.

Genomic distribution of ChIP-seq peaks from binding events for wild-type GL2 and mutant versions are indicated (FDR < 0.1). Most of the significant peaks occur in the intergenic and promoter regions. TSS, transcription start site; TTS, transcription termination site; UTR, untranslated region.

**Summary of GL2 binding sites in the *Arabidopsis thaliana* genome**

| **Sample** | | **Intergenic** | **Promoter-TSS** | **Exon** | **TTS** | **Intron** | **5'-UTR** | **3'-UTR** | **Total** |
|---|---|---|---|---|---|---|---|---|---|
| Wild type | *EYFP:GL2* | 306 | 246 | 122 | 160 | 83 | 7 | 1 | 925 |
| Mutants | *EYFP:gl2*$^{Amo}$ | 1298 | 782 | 668 | 502 | 262 | 10 | 11 | 3533 |
| | *EYFP:gl2*$^{\Delta ST}$ | 1018 | 564 | 442 | 320 | 177 | 4 | 6 | 2541 |
| | *EYFP:gl2*$^{E375G;R392M}$ | 419 | 344 | 186 | 195 | 128 | 10 | 2 | 1284 |
| | *EYFP:gl2*$^{\Delta HD6;\Delta ZLZ}$ | 0 | 0 | 2 | 0 | 0 | 0 | 0 | 2 |
| | *gl2-5* | 34 | 65 | 52 | 49 | 26 | 1 | 0 | 227 |

**Supplemental Table 3.** Oligonucleotides used in this study.

| **I. Primers for constructing mutations in *GL2*.** Nucleotide bases in red denote changes leading to amino acid subsitutions. | |
|---|---|
| **Name** | **5'-3' sequence** |
| gl2_K146T_F | CT CGC CAG GTC ACG TTC TGG TTC |
| gl2_K146E_F | CT CGC CAG GTC GAG TTC TGG TTC |
| gl2_K146T or E_R | G GGC CAG CCC TAG TTG CTT GC |
| gl2_ΔHD11 or 46_F | GCT ATT CAA GAA CGG CAC GAG AAC |
| gl2_ΔHD20_F | CAG CAG CTG AGC AAG CAA C |
| gl2_ΔHD20 or 46_R | ATC GGT GGT GTG ACG ATG ATA C |
| gl2_ΔHD11_R | CTT GAC CTG GCG AGG GG |
| GL2_START_ Δ _F | [Phos] GTC TTC TTC ATG GCT ACC AAC GTC CCC ACC (Schrick et al., 2014) |
| GL2_START_ Δ _F | [Phos] GAG GGC AAA GAC GCC CGT GTA GAA ATC G (Schrick et al., 2014) |
| GL2_HD_Δ_F | [Phos] GAA CGG CAC GAG AAC TCC CTG CTC |
| GL2_HD_Δ _R | [Phos] CTT ATT AGT GCC CTT GTT TCC AGC TGC G |
| GL2_ZLZ_Δ _F | [Phos] TCA TGC TCC GAC GAT CAA GAA CAC CG |
| GL2_HD-ZLZ_Δ _R | [Phos] TGT GCG GCG GTT TTG GAA CCA GAA CTT G |
| GL2_W279R_F | A GGC GAA CCT ATG AGG CTC CGC AGC G |
| GL2_W279R_R | GA GGT GGC CAT CTT CTG GAG TTC AAG GGT G |
| GL2_E375G_F | CAG CTG ATG TTC GGA **GGG** ATG CAG CTG CTC ACT |
| GL2_E375G_R | AGT GAG CAG CTG CAT **CCC** TCC GAA CAT CAG CTG |
| GL2_R392M_F | GAA GTG TAC TTC GTG **ATG** AGC TGC CGG CAG CTG |
| GL2_R392M_R | CAG CTG CCG GCA GCT **CAT** CAC GAA GTA CAC TTC |
| GL2_L480P_F | G GTC GCC ACC **CCT** CAG CTC CAT TGC G (Wojchiechowska et al, 2021) |
| GL2_L480P_R | C GCA ATG GAG CTG **AGG** GGT GGC GAC C (Wojchiechowska et al, 2021) |
| GL2_A258P_F | GAAGTCCCGTATT **CCC** GAGATTTCTAACCG |
| GL2_A258P_R | TCGAGGGCAAAGACGCCCGTGTAGAAATC |
| GL2_I289N_F | GACTGGCCGTGAG **AAT** CTCAACTACGATG |
| GL2_I289N_R | TCAACGCTGCGGAGCCACATAGGTTCG |
| GL2_E294Q_F | TCTCAACTACGAT **CAG** TACCTCAAGGAG |
| GL2_E294Q_R | ATCTCACGGCCAGTCTCAACGCTG |
| GL2_M271I_F | TGAACTCCAGAAG **ATT** GCCACCTCAGGC |
| GL2_M271I_R | AGGGTGGCTCGGTTAGAAATCTCGGC |
| **II. Sequencing primers for proGL2:EYFP:GL2 cassette** | |
| proGL2::EYFP:GL2_seq_B1 | CAC AGG AAA CAG CTA TGA CC |
| proGL2::EYFP:GL2_seq_B2 | CGA TCG GTC AAT GCC TCT CGC |
| proGL2::EYFP:GL2_seq_B3 | GAG TCC CAG TTT CCT ATA ATC C |
| proGL2::EYFP:GL2_seq_B4 | CCT TCC CCC CCT GTC TAC AG |
| proGL2::EYFP:GL2_seq_B5 | GTA TCA TTG GAA TTG TAG AGG C |
| proGL2::EYFP:GL2_seq_B6 | CAA GGA CGA CGG CAA CTA C |
| proGL2::EYFP:GL2_seq_B7 | GCC CTC TCT CTA TCT CTC GC |
| proGL2::EYFP:GL2_seq_B8 | GGC TAA TTC CTC CTG CCC C |
| proGL2::EYFP:GL2_seq_B9 | CGA TGT TAT CCG GCA AGG CG |
| proGL2::EYFP:GL2_seq_B9_5 | GTC AAC ACC GGT TTG GCC |
| proGL2::EYFP:GL2_seq_B10 | GCT TCT ACC GCG CCA TTG C |
| proGL2::EYFP:GL2_seq_B11 | CGT CTA CAC AAG ACG ACA G |
| proGL2::EYFP:GL2_seq_B12 | GGG TTT GCC TCC AAC CCC |
| proGL2::EYFP:GL2_seq_B13 | GG AGT GCG AGG AGA AGA GGG |
| proGL2::EYFP:GL2_seq_B14 | GTC GTG ATC CTC ACC CTC C |

| | |
|---|---|
| **III. Primers for amplification of *GL2* cDNA from pENTR/D-TOPO vector to clone into binary vector SR54 and for sequence verification.** | |
| EYFP:GL2_F | TCA AGC TTC GAA TTC TGC AGT CGA CAT GTC AAT GGC CGT CGA CAT G |
| p1300_GL2_R | CCA CCG CGG TGG AGC TCG TCA TTA GCA ATC TTC GAT TTG |
| **IV. Primers for genotyping mutants** | |
| To confirm *gl2-5* mutation | |
| GL2_F_112 | ATGTCAATGGCCGTCGACATGTC (Wang et al., 2007; Khosla et al., 2014) |
| En8130 | GAGCGTCGGTCCCCACACTTCTATAC (Baumann, 1998; Khosla et al., 2014) |
| To confirm transgene (*GL2* cDNA in *proGL2:EYFP:GL2*) | |
| SR54_GL2_747F | GTC GAC ATG TCA ATG GCC GTC G |
| SR54_GL2_747R | AGC TCG TCA TTA GCA ATC TTC GAT TTG |
| **V. Primers for constructing K107T, K107E and START deletion in PDF2 and ATML1.** Nucleotide bases in red denote changed bases underlying respective amino acid substitutions. | |
| pdf2_K107T_F | CT CTT CAA GTT AAG TTT TGG TTC C |
| pdf2_K107E_F | CT CTT CAA GTT GAG TTT TGG TTC C |
| pdf2_K107E_R | G CTC TAA ATT GAG ATC ACG GC |
| pdf2ΔSTART_F | GCT AGC TCC ATG GCC AGC |
| pdf2ΔSTART_R | AGA AGG AAT CGA AAC TGA CCT C |
| atml1ΔSTART_F | GCC AGT TCC ATG GCC AGC |
| atml1ΔSTART_R | AGA AGG TAT CGA AAC CGA CCT C |
| **VI. Primers for confirming Gateway cloning of pIX-HALO constructs** | |
| pIX-Halo_Seq | GCCTAACTGCAAGGCTGTGG |
| P1300_PDF2_R | ACCGCGGTGGAGCTCG CTA CGC TCC TCC TCC AAC |
| P1300_ATML1_R | CACCGCGGTGGAGCTCG TTA GGC TCC GTC GCA GGC C |
| **VII. Oligonucleotides used in electrophoretic mobility shift assay (EMSA).** Three repeats of the L1 motif are depicted in red. | |
| L1_EMSA_F_Cy3 | /5Cy3/GGG TACATTTA TACATTTA TACATTTA AAT |
| L1_EMSA_R | ATT TAAATGTA TAAATGTA TAAATGTA CCC |
| **VIII. Primers for restriction enzyme cloning into pGBK and pGAD vectors (Clontech Matchmaker)** | |
| GL2_747aa_EcoRI_F | GAC AGG **GAA TTC** *ATG* TCA ATG GCC GTC GAC ATG TCT TCC |
| GL2_747aa_BamHI_R | TGCTGT**GGATCCTCATCA** GCA ATC TTC GAT TTG TAG AC |
| GL2_493aa_BamHI_R | TTGGT**GGATCCCTA** GGT AGC CAT GAA GAA GAC AAG GCG |
| GL2_253aa_EcoRI_F | GTCTTT**GAATTC** GAG AAG TCC CGT ATT GCC GAG ATT TC |
| GL2_494aa_EcoRI_F | TTCATG**GAATTC** AAC GTC CCC ACC AAA GAC TCT CTC GG |
| GL2_157aa_EcoRI_F | ACCGCCGCACA**GAATTC** AAG GCT ATT CAA GAA CGG C |
| GL2_231aa_BamHI_R | ATCGTC**GGATCCTCA** AGC CTG CAG GGG ATA GGG |
| **IX. Primers for sequence verification of Y2H constructs** | |
| BD_seq_F | AACCGAAGTGCGCCAAGTGTCTG |
| AD_seq_F | TATAACGCGTTTGGAATCACT |
| BD & AD_seq_R | AGCCGACAACCTTCATTGGAGAC |
| **X. Primers for cloning in pENTR$^{TM}$/D-TOPO (Start and Stop codons, red)** | |
| GL2_TOPO_F | CACC **ATG** TCA ATG GCC GTC GAC ATG |
| GL2_TOPO_R | **TCA TTA** GCA ATC TTC GAT TTG TAG ACT TC |
| **XI. Primers for sequence verification of BiFC Gateway constructs** | |
| GFP-NXGW_seq | GACAAGCAGAAGAACGGCAT |
| pro35S_pK7WG2_seq | GGTGGCACCTACAAATGCCATC |

**Supplemental Data Set 1.** Mapping statistics of sequencing reads from ChIP-seq data.

**(See xls spreadsheet)**

The number of reads, alignment statistics, and total mapped reads to the *Arabidopsis thaliana* genome are indicated for individual input and immunoprecipitated (IP) samples. The total mapped reads of merged replicate IP samples are shown at the bottom. The ChIP-seq assays were performed with seedlings expressing EYFP-tagged GL2 wild-type and mutant proteins, including three START domain mutants (gl2_ΔST (listed here as gl2_dSTART), gl2_Amo, and gl2_E375G;R392M) as well as a mutant deleted for the six terminal amino acids of the HD and the entire leucine zipper (gl2_ΔHD6;ΔZLZ (listed here as gl2_dZip)).
